# TRPV4 interacts with mitochondrial proteins and acts as a mitochondrial structure-function regulator

**DOI:** 10.1101/330993

**Authors:** Ashutosh Kumar, Rakesh Kumar Majhi, Tusar Kanta Acharya, Karl-Heinz Smalla, Eckart D. Gundelfinger, Chandan Goswami

**Affiliations:** National Institute of Science Education and Research Bhubaneswar, School of Biological Sciences, P.O. Jatni, Khurda 752050, Odisha, India; Homi Bhabha National Institute, Training School Complex, Anushakti Nagar, Mumbai 400094, India; Leibniz Institute for Neurobiology and Center for Behavioral Brain Sciences (CBBS), Brenneckestr 6, Magdeburg 39118, Germany; Medical Faculty, Otto von Guericke University, Magdeburg, Germany

**Keywords:** TRP channels, Mitochondrial dynamics, Mitochondrial calcium, Metabolism, Mitochondrial diseases

## Abstract

TRPV4 has been linked with the development of sensory defects, neuropathic pain, neurodegenerative disorders such as Charcot Marie Tooth disease and various muscular dystrophies. In all these cases mitochondrial abnormalities were tagged as cellular hallmarks and such abnormalities have been reported as key factor for the pathophysiological conditions. Mitochondria also have the unique ability to sense and regulate their own temperature. Here, we demonstrate that TRPV4, a thermosensitive ion channels, localizes to a subpopulation of mitochondria in various cell lines, in primary cells and also in sperm cells. Improper expression and/or function of TRPV4 induce several mitochondrial abnormalities such as low oxidative potential, high Ca^2+^-influx and changes in electron transport chain functions. TRPV4 is also involved in regulation of mitochondrial morphology, smoothness, and fusion-fission events. The C-terminal cytoplasmic region of TRPV4 can localize it to mitochondria and interacts with mitochondrial proteins including Hsp60, Mfn1 and Mfn2. Regulation of mitochondria by TRPV4 may contribute to previously uncharacterized mitochondria-specific functions observed in various cell types. This discovery may help to link TRPV4-mediated channelopathies with mitochondria-mediated diseases.

**Abbreviations**
ER: Endoplasmic Reticulum
IMM: Inner Mitochondrial Membrane
MAM: Mitochondria-associated ER membrane
MCU: Mitochondrial uniporters
MOM: Mitochondrial outer membrane
NO: Nitric Oxide
ROS: Reactive Oxygen Species
TRPV4: Transient Receptor Potential Channel Vanilloid subfamily member 4
VDAC: Voltage Gated Anion Channels

## Introduction

It is well established that intracellular Ca^2+^-homeostasis is maintained by Ca^2+^-binding proteins present in cytoplasm, endoplasmic reticulum (ER) and mitochondria (Kostyuk and Verkhratsky. 1994; Svichar et al. 1997; Verkhratsky et al. 1998). The importance of the later organelle is not fully understood. Mitochondria are important cellular organelles, which can modulate both the amplitude and the spatio-temporal patterns of Ca^2+^-signals (Budd and Nicholls. 1996; Duchen et al. 1999; Jouaville et al. 1995). However, the array of mitochondrial proteins and/or ion channels involved in maintenance of mitochondrial Ca^2+^-homeostasis and the actual mechanisms involved within remain as enigma. It has been demonstrated that the distribution of mitochondria is particularly dense in the areas where cytoplasmic Ca^2+^ concentration is high, and such heterogeneous distribution correlates well with the distribution of Ca^2+^-selective ion channels present in ER (such as Inositol trisphosphate receptor), sarcoplasmic reticulum (such as Ryanodine receptor) and in plasma membrane (such as Voltage-operated channels and Store-operated channels) (Rizzuto al et al. 1998; Csordas et al. 1999; Szalai et al. 2000; Mannella et al. 1998). This implies that mitochondria are often closely associated with these microdomains of high cytoplasmic Ca^2+^, which facilitate influx and efflux of Ca^2+^ across the mitochondrial membrane. Because of this notion many mitochondria show close proximity with the ER (commonly known as ER-associated mitochondria) and mediate easy exchange of divalent cations such as Ca^2+^. Accordingly mitochondrial Ca^2+^-uptake is a key regulator for cell survival and various functions such as metabolism, secretion and cellular signalling. Within the cell, mitochondria maintain a higher temperature due to their unique ability to sense different temperatures and respond to slight changes in physiological temperatures (Chrétien et al. 2018; Lane N 2018). However, candidate proteins present in mitochondria that are involved mediating such mitochondrial functions, including Ca^2+^-transport or temperature sensing, have not been identified yet.

The permeability of Ca^2+^ in mitochondrial outer membrane (MOM) is known to be governed by various isoforms of Voltage Gated Anion Channels (VDAC) (Gincel et al. 2002). Therefore, Ca^2+^- uptake from the cytoplasmic microdomains becomes a limiting factor if the expression as well as activity of VDACs is low. Once free Ca^2+^ ion crosses the MOM, it further crosses the inner mitochondrial membrane (IMM) with the help of mitochondrial uniporters (MCU) abundantly present there (Kirichok et al. 2004). Though the molecular identities of these different MCUs are not clear, it is known that these MCUs are of low conductance with high selectivity for Ca^2+^. It has been reported that the MCU-mediated Ca^2+^-current could be inhibited by ruthenium red and gadolinium suggesting that channels inhibited by these agents are present there (Bernardi et al. 1984).

TRPV4 acts as a Ca^2+^-permeable nonselective cation channel and can be activated by different physical stimuli such as moderate temperature (> 27°C), low pH, change in osmolarity, mechanical force and by several chemical stimuli such as phorbol ester derivative 4α-phorbol 12, 13-didecanoate (4α-PDD), endocannabinoids and arachidonic acid (AA) metabolites (Guler et al. 2002; Watanabe et al. 2002; Watanabe et al. 2003; Suzuki et al. 2003; Strotmann et al. 2000; Liedtke et al. 2000). Apart from the peripheral neurons, expression of TRPV4 is also reported in different types of tissues and cells such as in lung, kidney, heart, gut, sensory neurons, sympathetic neurons, vascular smooth muscle cells, bone cells and endothelial cells (Chung et al. 2003; Fernandez-Fernandez et al. 2002; Jia et al. 2002; Liedtke et al. 2000; Strotmann et al. 2000; Wissenbach et al. 2000). In agreement with the involvement of TRPV4 in different cellular functions and expression of TRPV4 in these tissues, TRPV4 has been linked with different diseases and pathophysiological conditions (Verma et al. 2010; Nillus et al. 2013). In this work we demonstrate that the mitochondrial structure and dynamics can be regulated by TRPV4. We show that TRPV4 localizes in mitochondria and interacts with several key mitochondrial proteins, such as Hsp60, Mfn1 and Mfn2, which regulate mitochondrial structure and function. Furthermore, we demonstrate that TRPV4-mediated mitochondrial regulation is a common phenomenon observed in several cellular systems including neurons and in vertebrate spermatozoa suggesting that such regulation can be ubiquitous in nature.

## Materials and Methods

### Reagents and antibodies

Affinity-purified rabbit polyclonal antibodies were purchased from Alomone (Jerusalem, Israel) and Sigma-Aldrich (Bangalore, India) respectively. The TRPV4-specific peptide (CDGHQQGYAPKWRAEDAPL, used as a blocking peptide of the TRPV4 antibody) was purchased from Alomone (Jerusalem, Israel). Mouse monoclonal anti-Hsp60, anti-Mfn2, anti-Mfn1, anti DRP1, anti-Opa1, anti-Cyt C were purchased from Abcam (USA). Hsp60 and Cyt C antibodies were obtained from Santa Cruz Biotechnology (kind gift from Ansgar Santel, Berlin, Germany). Mouse and rabbit specific secondary antibodies coupled to Alexa fluor488 and Alexa fluor594 as well as specific dyes such as MitoTracker Red, Lysotracker Red, ER-tracker were purchased from Molecular Probes (Invitrogen, Bangalore). All other drugs namely 4αPDD, RN1734, Ionomycin and all mitochondrial complex chain inhibitors were purchased from Sigma Aldrich.

### Constructs and vector used

For expression of TRPV4 without any tag, full-length TRPV4 (NCBI accession number AF263521) cloned in pCDNA3.1 vector (a kind gift from Prof. Jon D Levine) was used as described before (Goswami et al. 2010). Similarly, for expression of TRPV4 as a GFP-tagged protein, TRPV4-GFP construct (cloned in pEGFPN3 vector, kindly provided by Prof. J. Berreiter-Hahn) was used (Becker et al. 2005). The transmembrane fragment of hTRPV4 (466-711 aa) was amplified by using 5`GTTGAATTCTTCGGGGCCGTCTCCTTCTAC3`and 5`GCCGTCGACTTAGAGGAGCAGCACAAAGGTGAG3` primer sequences respectively. This fragment is cloned into the pSEGFP vector (Addgene) at BamHI and SalI site and confirmed by sequencing. This fusion protein is termed as TRPV4-TM-GFP. The N-terminus fragment (aa 1-465) of Rat TRPV4 was amplified by using following primer sequences 5`CCGCTCGAGCTATGGCAGATCCTGGTGATGT3` and 5`CGCGGATTCCTAACGCCACTTGTCCCTCA3` primer sequences. Similarly, C-terminus fragment (aa 718-871) of Rat TRPV4 was amplified by using 5`CCGCTCGAGCTATGGGTGAGACCGTGGGCCA3`and 5`CGCGGATTCCTACAGTGGTGCGTCCTCCG3` primer sequences. The amplified DNA fragments were cloned into the RFP-vector (Clontech) at XhoI and BamHI site and confirmed by sequencing. These RFP-tagged constructs are termed as TRPV4-Nt-RFP, TRPV4-Ct-RFP respectively.

### Isolation of mitochondria from mammalian brain and CHO-K1 cells

Adult goat brains were obtained from a local slaughterhouse. The meninges were separated from the brain and the tissues were taken to the laboratory in isotonic mitochondrial isolating buffer (10 mM HEPES, 1 mM EDTA, 320 mM sucrose, pH 7.4) containing complete protease inhibitor cocktail. Isolation of mitochondria was based on the method established previously with some minor modifications (Whittaker et al. 1968; Frezza et al. 2007). Briefly, goat brain was homogenized in mitochondrial isolating buffer. After homogenization, the homogenate was centrifuged at 1000g for 10 minutes. The supernatant was collected and centrifuged at 10,000g for 20 minutes to get crude mitochondrial pellet fraction. The pellet fraction was washed twice with same buffer and centrifuged again at 10,000g for 20 minutes. The mitochondrial fraction was isolated as pellet. For isolation of mitochondria from CHOK1 cells, 90% confluent cells were seeded in 100 mm dishes and grown for 24 hours. Mitochondrial fraction and cytoplasmic fraction from the cells were isolated by using a mitochondrial isolation kit according to the manufacturer’s protocol (Sigma Aldrich, Bangalore).

### Interaction of TRPV4 fragments with isolated intact mitochondria

The C-terminus of TRPV4 was expressed as a MBP fusion protein (termed as MBP-TRPV4-Ct). Freshly isolated mitochondria (30 µg) from brain tissue were incubated with equal amount of purified MBP-TRPV4-Ct and MBP-LacZ at 25°C in the same mitochondrial isolating buffer in presence or in absence of Ca^2+^ (1 mM) or a mixture of GTP and ATP (1 mM each). After 30 minutes of incubation, mitochondria were washed two times (centrifuged at 5400 g for 5 minutes) with mitochondrial isolating buffer which separates the unbound MBP-TRPV4-Ct and MBP-LacZ from the mixture. The mitochondrial pellet was suspended in PEMS buffer (50 mM PIPES pH 6.8, 1 mM EGTA, 0.2 mM MgCl_2_, 150 mM NaCl) and gel sample was prepared. Complete protease inhibitor cocktail was added in each fraction and the sample was were boiled for 5 minutes in with (1x) Laemmli sample loading buffer for SDS-PAGE.

### MBP-pulldown assay for identifying TRPV4 interacting proteins

Expression and purification of the C-terminal cytoplasmic domain of TRPV4 fused with MBP (MBP-TRPV4-Ct) as well as LacZ fused with MBP (MBP-LacZ) were performed according to the protocol described previously (Goswami et al. 2004). In brief, MBP-TRPV4-Ct and MBP-LacZ were expressed in *E. coli* and the cleared cell lysates were applied to amylose resin (NEB) and incubated for 2 hours at RT. Approximately 50 µL of amylose resin per tube with the bound fusion protein was used for pull-down experiments. Depending on the respective experiments, the resin with coupled fusion protein was incubated with 60 µL of mitochondrial protein (0.7 mg/ml protein) for 2 hours at 4°C in the presence or absence of Ca^2+^ (1 mM) or a mixture of GTP and ATP (1 mM each). The amylose resin present in each tube was washed three times with PEM-S buffer. The proteins were eluted by using elution buffer (PEM-S supplemented with 20 mM maltose). Eluted samples were analysed by 10-12% SDS–PAGE and Western Blot analysis with candidate specific antibodies were performed.

### His-tagged pull down assay for Mfn1 and Mfn2 with MBP-TRPV4-Ct

His-tagged Mfn1 and His-tagged Mfn2 constructs were received as a gift from Prof. Naotada Ishihara (Institute of Life Science, Kurume University) (Ishihara et al. 2004). His-Mfn1 and His-Mfn2 constructs were freshly transformed into *E. coli* and protein was expressed in 2XYT media. Cells were suspended in lysis buffer (20 mM HEPES, pH 7.4, 250 mM NaCl, 1 mM PMSF and EDTA-free protease inhibitor) for 2 hours on ice and subsequently cells were disrupted by sonication (60 Hz, 10 cycles with 5 sec pulse interval) in ice cold condition. The sonicated lysate was then centrifuged at 10000 g for 2 hours and the supernatant was discarded as Mfn2 and Mfn1 appeared inside inclusion bodies. The pellet was taken in 30 ml lysis buffer and was resuspended by sonication. This was followed by the addition of 1 ml 20% Triton-X-100 and mixed well at 4°C for 30 minutes. This suspension was centrifuged at 10000 g for 40 minutes and the collected supernatant was incubated with Ni-NTA beads (pre-washed with pre-equilibration buffer: 20 mM HEPES, pH 7.4, 250 mM NaCl, 10 mM imidazole) for 2 hours at 4°C. It was then washed thrice with washing buffer (20 mM HEPES, pH 7.4, 250 mM NaCl, 20 mM imidazole) and then centrifuged at 400 g for 3 minutes. Subsequently, equal amount of MBP-TRPV4-Ct or MBP-LacZ were incubated with His-Mfn2 or His-Mfn1 coupled Ni-NTA resin in presence or absence of 1 mM Ca^2+^. Finally the proteins were eluted in 100 mM imidazole solution.

### Western blot analysis

To perform Western Blot analysis, protein samples were separated by SDS-PAGE (10-12%) and subsequently transferred to a PVDF membrane (Millipore) by semidry electroblotting (Bio-Rad). The membranes were blocked with 5% non-fat milk in TBS-T (20 mM Tris, 150 mM NaCl, 0.1% Tween-20) buffer followed by incubation with the primary antibodies overnight at 4°C. Membranes were washed in TBST buffer 3 times followed by incubation with appropriate HRP-conjugated secondary antibody (1:10000 dilution) (Amersham/BD life science). The specific protein signals were identified in Chemidoc (Bio-Rad) by using the SuperSignal West Femto Chemiluminescent reagent (Pierce). The dilutions of these primary antibodies were as follows: TRPV4 (1:200), Mfn1 (1:250), Mfn2 (1:250), Cyt-C (1:250), Hsp60 (1:500), Opa1 (1:250), MBP (1:20000), Tubulin (1:500). For peptide blocking experiments (against rabbit polyclonal anti-TRPV4), a peptide (CDGHQQGYAPKWRAEDAPL) corresponding to amino acid residues 853-871 of rat TRPV4 (NCBI Accession no Q9ERZ8) was used to confirm the specificity of the immunoreactivity. The blocking peptides were used at 1:3 dilutions.

### Labelling of mitochondria with MitoTracker Red

Cells were grown and transfected on 12 mm glass cover slips. Two days after seeding or transfection, cells were incubated with MitoTracker Red (1 μM) for 20 minutes in cell culture incubator. Subsequently cells were washed with 1 X PBS and fixed by 4% PFA at RT. Similarly for sperm cells (floating cell) after drug treatment, MitoTracker Red (2 μM) was added in sperm incubating media and kept at 37°C in water bath for 20 minutes. Subsequently incubated cells were diluted in 3 ml of 1 X PBS to avoid any clumping or aggregation during PFA fixation and immediately equal volume of 3 ml PFA (4% PFA) were added in diluted sperm for fixation.

### Mitochondrial Ca^2+^ imaging

Mitochondrial Ca^2+^ imaging was performed with ratiometric mitochondrial pericam constructs (A gift from Dr. Atsushi Miyawaki, Saitama, Japan). Ratiometric pericam is a Ca^2+^-indicator fusion of the yellow fluorescent protein (YFP) and Calmodulin which is able to enter mitochondria. Binding of Ca^2+^ to Mito-pericam changes its excitation wavelengths from 415 nm to 494 nm while its emission spectrum is maintained at 515 nm. Mito-pericam was expressed in CHOK1-V4 and CHOK1-Mock cells by transient transfection and 24 hours after transfection, cells were imaged with confocal microscopy.

### JC-1 (Ratiometric dye) staining in adherent cells

Mitochondrial potential staining in adherent cell (CHOK1-V4 and CHOK1-Mock) were performed with 5,5,6,6-tetrachloro-1,1,3,3-tetraethylbenzimidazolylcarbocyanine iodide (JC-1) cationic dye. JC-1 dye exhibits potential-dependent accumulation inside the mitochondria and also has fluorescence emission properties that shift from green (525 nm) to red (590 nm) region (Reers et al. 1995). TRPV4 specific activator 4αPDD (5 μM) and inhibitor RN1734 (10 μM) were added for 6 hours and after that JC-1 (5 µM) was added for 40 minutes and kept in the incubator. Subsequently cells were washed with 1 X PBS and coverslips were taken for live cell imaging by confocal microscopy (Zeiss LSM780).

### Immunocytochemistry, live cell imaging and microscopy

Cells were grown and transfected on 12 mm glass cover slips. Two days after seeding or transfection, cells were fixed either with 4% paraformaldehyde at room temperature (RT). Cells were permeabilized with 0.1% Triton X 100 in PBS for 5 minutes, followed by two times washing with 0.1% PBS-T (0.1% Tween 20 in 1 X PBS). Cells were blocked with 5% bovine serum albumin (BSA) in PBS-T. Thereafter, cells were incubated with different primary antibodies such as Hsp60 (1:800), Cyt C (1:250), TRPV4 (1:500) for overnight. Thereafter cells were washed three times with PBS-T (PBS supplemented with 0.1% Tween 20). The cells were further incubated for 1 hour with Alexa dye labelled secondary antibody (anti-mouse or anti-rabbit) diluted (1:1000) in PBS-T and 5% bovine serum albumin (BSA) in 1:1 ratio. Cells were incubated with DAPI (5 μM) in PBS-T for 30 minutes at RT. Subsequently cells were washed with PBS two times and cover slips were finally mounted onto glass slides with Fluoromount G (Southern Biotech). The cells were imaged by using a confocal microscope (Zeiss, LSM780) using the 63X objective. All images were taken under same conditions for comparative analysis. Images were analyzed and processed by using LSM Image examiner software and by ImageJ (Mitochondrial morphology plugin).

### Assay for mitochondrial electron transport chain and MPT

All enzymatic activity was determined in freshly isolated goat brain mitochondria and performed at 30°C in assay buffer (1 ml) consists of 20 mM potassium phosphate buffer (pH 7.2), EDTA (0.1 Mm), MgCl_2_ (10 mM), mitochondria (30 µg). Before starting the enzymatic assay, isolated mitochondria were pre-incubated with TRPV4-specific activator 4αPDD (5 µM) or inhibitor RN1734 (20 µM). For measuring complex I activity, NADH-based spectroscopic analysis was performed with some minor modification (Kramer et al. 2005; Humphries et al. 1998). The absorbance of reduced DCPIP (100 µM) was measured in presence of NADH (50 µM) and decylubiquinone (150 µM) at 600 nm. Complex II activity was measured at 600 nm using DCPIP (50 µM) as electron acceptor from artificial substrate decylubiquinone (DU, 50 µM), as it accepts electron from complex II and transfer to DCPIP. Complex III activity was measured by monitoring the reduction of cytochrome C (100 µM) after adding of reduced decylubiquinone (DUH_2_, 100 µM) in assay buffer at 550-540 nm. Complex IV activity was measured by the oxidation of reduced cytochrome C (50 µM) in presence 20 µg mitochondrial protein at 550 nm.

Mitochondrial swelling or membrane permeability transition pore induced by influx of excess Ca^2+^ into mitochondria which can be detected as a decrease in light scattering property in isolated mitochondria at 540 nm (Kobayashi et al., 2003). Briefly, Mitochondria (100 μg) were treated with TRPV4 activator and inhibitor for 15 minutes in mitochondrial isolating buffer. Subsequently, treated mitochondria were transferred to mitochondrial swelling buffer [HEPES (20 mM), KH_2_PO_4_ (2 mM), KCl (125 mM), EGTA (1 μM), MgCl_2_ (1 mM), Malate (5 mM), Glutamate (5 mM), pH: 7.4] and decay in absorbance was monitored at 540 nm.

### Cell culture, transfection and pharmacological modulation of TRPV4

Chinese hamster ovary (CHOK1) cells stably selected for TRPV4 (cloned in pCDNA3.1 vector, termed as CHOK1-V4 cells) or empty pCDNA3.1 vector only (termed as CHOK1-Mock cells) were grown in F-12 Ham’s medium containing FBS (10% v/v), L-glutamine (2 mM), Streptomycin (100 µg/ml), Penicillin (100 U/ml), in a humidity controlled incubator maintained with 5% CO_2_ and at 37°C. HaCaT cells were cultured in DMEM media supplemented with FBS (10% v/v), L-glutamine (2 mM), Streptomycin (100 µg/ml), Penicillin (100 U/ml), in a humidity controlled incubator at 37°C. Similarly, Human Umbilical Vein Endothelial cells (HUVEC) were cultured in endothelial cell growth media (EGM, Lonza) containing heat-inactivated Fetal Bovine Serum (FBS, 2%), human VEGF, epidermal growth factor (EGF), insulin-like growth factor-1 (IGF-1), basic fibroblast growth factor (FGFB) and amphotericin-B. Transient transfection was performed by Lipofectamine 3000 Plus reagent (Invitrogen) according to the manufacturers protocol. Generally 24-36 hours after transfection, the cells were used for live cell imaging or immunocytochemistry. For activation or inhibition of TRPV4, cells were either treated with TRPV4 activator (4αPDD) or inhibitor (RN1734) for the desired time. Subsequently, cells were either washed and processed for live cell imaging or fixed with 4% PFA.

### Collection of sperm cells

The mature sperm cells from fish was collected as mentioned before (a kind gift from CIFA, Bhubaneswar) (Majhi et al. 2013). The bull sperm was collected as described before (Approval number, NISER-IAEC/SBS-AH/07/13/10) (Kumar et al. 2016). The human sperm was collected as per the IEAC approval (NISER/IEAC/2015-11) following the process as described before (Kumar et al. 2016). In all cases, cells were fixed with PFA and in some cases labelled with MitoTracker-Red before these samples were fixed.

### Fractionation of inner and outer mitochondrial membrane

The inner and outer mitochondrial membranes were fractionated by using Digitonin with minor modification (Nishimura et al. 2014; Nishimura and Yano.2014; Pallotti et al. 2007; Schnaitman et al. 1968). Upon treating mitochondria with Digitonin the outer membrane along with the inter membrane space proteins get separated from the mitoplast (inner mitochondrial membrane and mitochondrial matrix). Freshly isolated mitochondrial pellets were treated with 1 ml of Digitonin buffer (MT isolation buffer containing 10mg/ml Digitonin). The suspension was mixed properly by using vortex mixer. After 30 minutes of incubation, the suspension was centrifuged at 12000g (20 min, 4°C). The pellet contains mitoplast (inner membrane plus matrix) and the supernatant contains solubilised outer membrane and inter-membrane space proteins. The supernatant was carefully collected and the pellet was resuspended in mitochondria isolation buffer. The mitochondrial suspensions were immediately boiled for 5 minutes in sample loading buffer for SDS-PAGE and Western Blot applications.

### Staining and imaging of isolated mitochondria obtained from brain tissue

Freshly isolated mitochondrial pellet from mouse brain tissue was re-suspended with mitochondria isolation buffer. The solution was treated with 1µm MitoTracker red and mixed thoroughly. After 30 minutes of incubation at 37°C, the mitochondria were fixed with 4% PFA.

TRPV4 was probed with primary antibody supplied by Alomone lab and detected by AlexaFlour488. The Hsp60 was probed with primary antibody supplied by Abcam and was detected by AlexaFlour594. Isolated mitochondria were imaged in Zeiss LSM 880 confocal microscope in Airyscan mode to obtain high resolution images of Mitochondria and TRPV4. Synaptic fractions were prepared according to protocols described previously (Goswami et al. 2010).

## Results

### TRPV4 is localized in mitochondria

Previously we have reported that in case of over expression, TRPV4 as well as TRPV4-GFP effectively localize in plasma membrane in F11 cells, a peripheral dorsal root ganglion (DRG) neuron-derived cell line. In addition a considerable fraction of TRPV4 localizes to intracellular organelles (Goswami et al. 2010). In order to characterize this intracellular fraction in more detail, we expressed TRPV4 in several neuronal as well as non-neuronal cell lines, i.e., in F11, COS7, HaCaT and HeLa cells, and performed immunolocalization studies. In all cases, we observed TRPV4 in the plasma membrane and to a lower extent in the cytoplasm. However, we noted that in a large number of cells TRPV4 is present in separate intracellular organelles (Supplementary figure 1A). Furthermore, TRPV4 immunostaining was performed in CHOK1 cells that are stably selected for TRPV4 (named as CHOK1-V4) and mock plasmid (named as CHOK1-Mock). Finally, to exclude that such localization is due to over-expression, endogenous TRPV4 was detected in HUVEC (primary cells). Very similar perinuclear localization of TRPV4 was observed in CHOK1-V4 cells and HUVEC cells (Supplementary figure 1B). In general, these organelle structures are located mainly at the perinuclear region and are mostly short cylindrical or spherical in nature. In most cases, these structures are straight without any bends or kinks.

To characterize these TRPV4-positive subcellular organelles in detail, we used HaCaT cell-line. First, we performed co-localization experiments with different subcellular organelle specific markers. Co-localization of TRPV4-GFP with MitoTracker Red indicated that TRPV4 may occur in mitochondria (Fig 1a). To confirm this we show that TRPV4 co-localizes with two endogenous mitochondrial markers, i.e. Cyt C and Hsp60, within mitochondrial structures (Fig 1b-c). TRPV4 seems to be present in some, but not in all mitochondria, indicating that the presence of TRPV4 in mitochondria is heterogeneous in nature. To rule out the possibility of co-localization as fixation artefact, we performed live cell imaging. We observed that in living HaCaT cells, TRPV4-GFP perfectly co-localizes with mito-RFP or mitoDsRed even under low expression conditions (Fig 1d-e). To explore if TRPV4 localizes in sub-cellular organelles other than mitochondria; HaCaT cells were transfected with TRPV4-GFP alone or with either ER-CFP, or Golgi-CFP, or peroxisome-CFP. These experiments suggest that TRPV4 does not occur at significant levels in ER or Golgi structures or peroxisomes (Supplementary figure 2). In certain cases, limited co-localization was observed with TRPV4 and peroxisome-CFP. Basically no co-localization was observed when TRPV4-GFP expressing HaCaT cells were immunostained with specific antibodies detecting ER (immunolabelled by KDEL ab), Golgi (labelled by Calnexin ab) or lysosomal structures (labelled by Lysotraker Red) (Supplementary figure 2).

**Figure 1.**
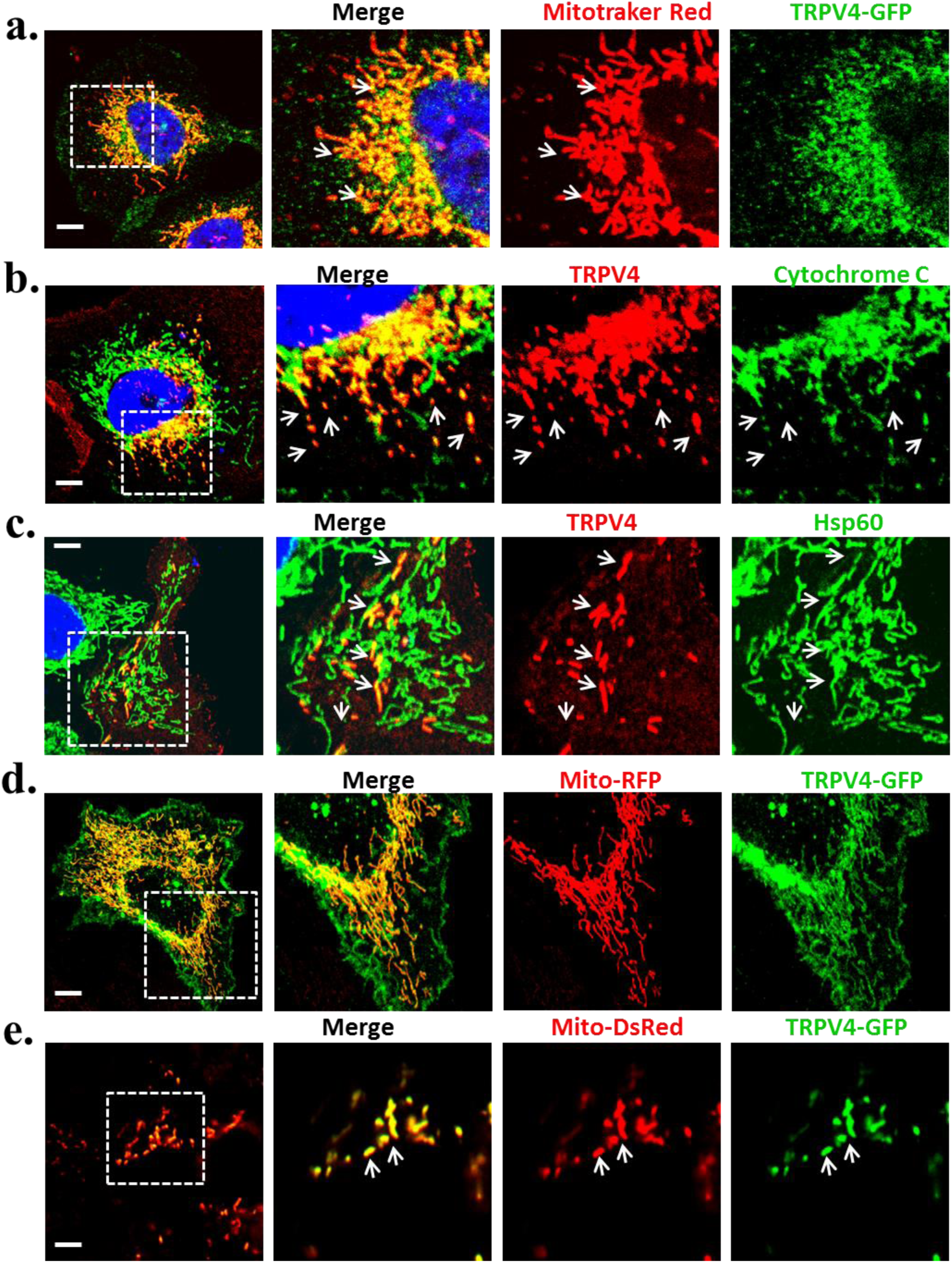
TRPV4 colocalizes with mitochondrial markers. **a.** Confocal images of HaCaT cells transiently expressing TRPV4-GFP and labelled with MitoTracker-Red. **b-c.** HaCaT cells transiently expressing TRPV4 immunostained with specific antibodies detecting Cytochrome C (b) and Hsp60 (c) are shown. TRPV4 colocalizes with these markers in a subset (indicated by arrows) but not in all mitochondria. **d.** Cells co-expressing Mito-RFP and TRPV4-GFP shows colocalization of TRPV4 with MitoDsRed. **e.** Confocal images of live HaCaT cells transiently expressing TRPV4-GFP and mitoDsRed. Enlarged confocal images depicting that in case of low expressing TRPV4-GFP positive cells, TRPV4 perfectly colocalizes with MitoDsRed within the mitochondria (indicated by arrows).

### TRPV4 is present in mitochondria endogenously

Next, we sought to confirm that TRPV4 is endogenously present in mitochondria. To this end, freshly isolated mitochondrial fractions from rat brain were labelled with mitochondria-specific dye MitoTracker Red. in *ex vivo*, After fixation they were stained for TRPV4 and analysed by super resolution microscopy (Fig 2a). In a same manner, isolated mitochondria were probed for both Hsp60 and TRPV4 (Fig 2a). In both cases TRPV4 was detected in a subset of mitochondria, but not in all. Next we tested the presence of TRPV4 in the mitochondria by immunoblot analysis. TRPV4 is present and enriched in the isolated mitochondria from rat brain (Fig 2b). TRPV4 is also detected and even enriched in mitochondria isolated from white adipose tissue (goat) and from CHOK1-V4 stable cell lines. In these fractions, we could detect TRPV4 as a band of ∼98 kDa, the expected size of the full-length TRPV4. Notably, TRPV4 is not detected in the mitochondria isolated from CHOKI-Mock cells. The TRPV4 specific immunoreactivities can be effectively blocked by the presence of a specific peptide (corresponding to the C-terminal sequence of TRPV4), suggesting that these bands are specific for TRPV4 (Fig 2c).

**Figure 2.**
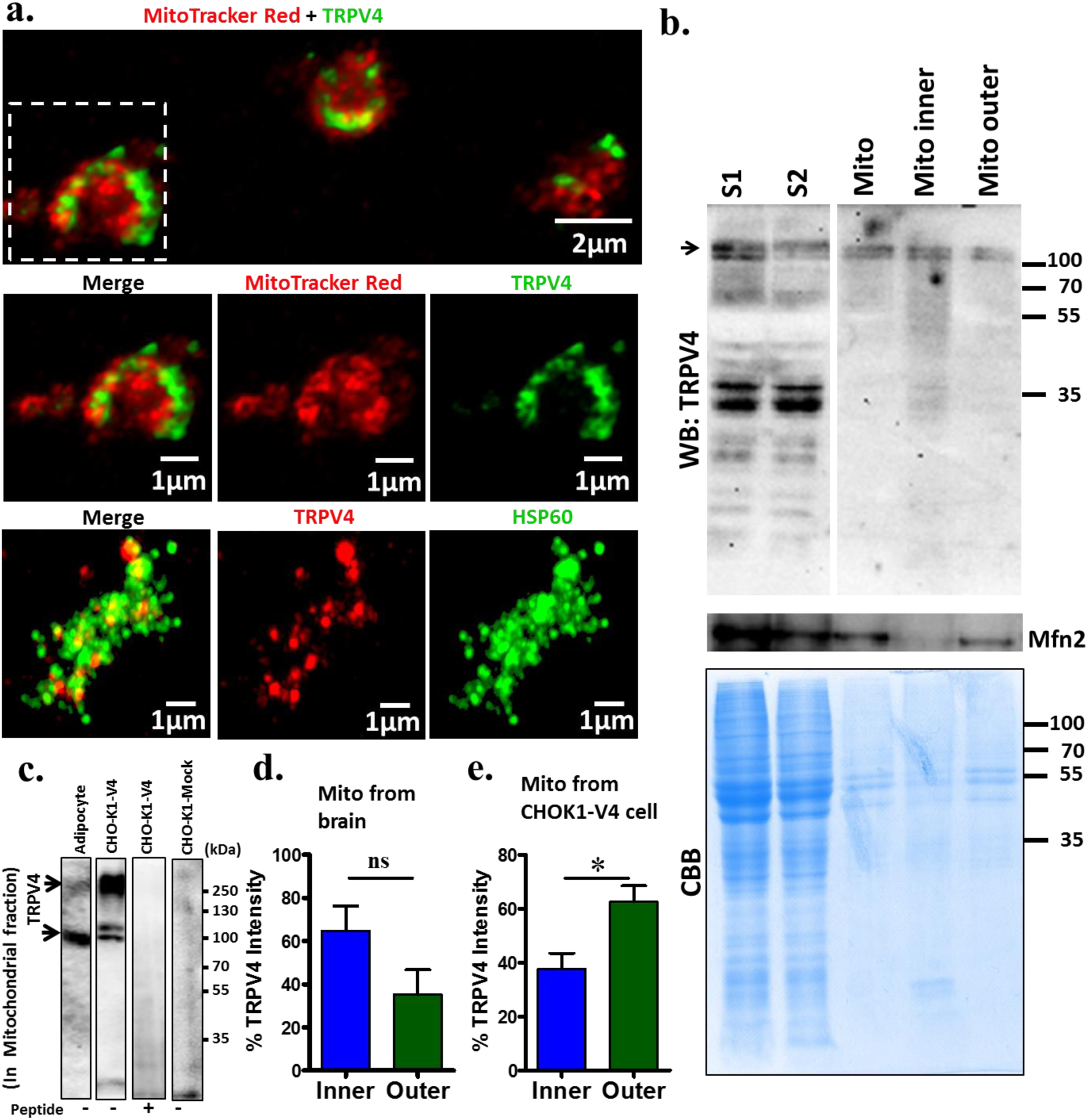
TRPV4 is endogenously present in mitochondria. **a.** Shown are the super resolution images of mitochondria isolated from rat brain and immunostained for TRPV4 (green). Isolated mitochondria were labelled with MitoTracker Red (upper and middle panel) or immunostained for Hsp60 (green, lower panel). TRPV4 is present in a subset of mitochondria, but not in all. **b.** Western blot analysis of different mitochondrial fractions (S1, S2 and Mitochondrial fraction) isolated from rat brain are shown. Endogenous TRPV4 is present in isolated mitochondrial fraction as well as in both inner and outer membrane fractions. **c.** Western blot analysis of mitochondrial fraction isolated from adipocytes, ChoKI-TRPV4 cells and CHO-KI-Mock cells probed with anti TRPV4 antibody in absence or presence of specific blocking peptide is shown. Immunoreactivity at the specific size (100 kDa) or at higher molecular weight is indicated by arrows. **d-e.** Quantification of relative abundance of TRPV4 in outer and inner mitochondrial membrane is shown (total mitochondrial TRPV4 is considered as 100%). Majority of the endogenous TRPV4 is present in the inner membrane rather than in outer membrane of mitochondria isolated from brain (ns = non-significant, n = 4, P value = 0.1181). In contrast, majority of the TRPV4 is present in the outer membrane rather than in the inner membrane of the mitochondria isolated from CHOKI-TRPV4 cell (n = 6, P value = 0.0141, n = 6).

Synaptic junctions present in neuronal tissues are highly energy demanding and accordingly they accumulate mitochondria. This is also reflected in isolated synaptosomes. To explore the presence of endogenous TRPV4 in synaptosomal protein fractions, density gradient fractionation was performed with rat forebrain homogenate. Different fractions were probed for the presence of TRPV4 with specific antibody. Immunoblot analysis revealed TRPV4-specific bands in the light membrane fraction and in the synaptosomal fraction, but not in the postsynaptic density fraction which contains primarily synaptic junctional proteins (Supplementary fig 3).

Next we attempted to explore if TRPV4 is present in the inner or outer membrane of mitochondria. For that purpose we isolated inner membrane and outer membrane and analysed the presence of TRPV4 there. We noted that TRPV4 is present in both inner and outer membrane when isolated from rat brain, where tentatively more TRPV4 seems to be present in the inner membrane. However in case of mitochondria isolated from CHOKI-V4 cells, TRPV4 seems more abundant in outer membrane than in the inner membrane (Fig 2d-e).

### TRPV4-positive mitochondria display an altered mitochondrial morphology

To explore the effect of TRPV4 activator 4αPDD or inhibitor RN1734 upon mitochondrial morphology, CHOK1-V4 and CHOK1-Mock cells were transiently transfected with mitoDsRed and subsequently treated with TRPV4 activator or inhibitor. Results indicate that in CHOK1-V4 cell, mitochondrial morphology is mainly spherical or round ball shaped (Fig 3a). In CHOK1-Mock cells mitochondrial morphology was normally elongated and tubular in shape even when cells were treated with TRPV4 activator or inhibitor for the same duration (Fig 3b). However in CHOK1-V4 cell, after the treatment of TRPV4 activator 4αPDD, most of the spherical shape (small) mitochondria formed aggregates (Represented with arrow in right side, Fig 3a,c). These aggregated mitochondria co-localize with TRPV4. In contrast, in the presence of TRPV4 inhibitor RN1734 mitochondrial morphology did not change much. CHOK1-Mock cells do not show significant change in presence of TRPV4 activator or inhibitor (Fig 3b,c). This indicates that activation of TRPV4 causes aggregation of mitochondria and increases their perimeter as well as their area.

**Figure 3.**
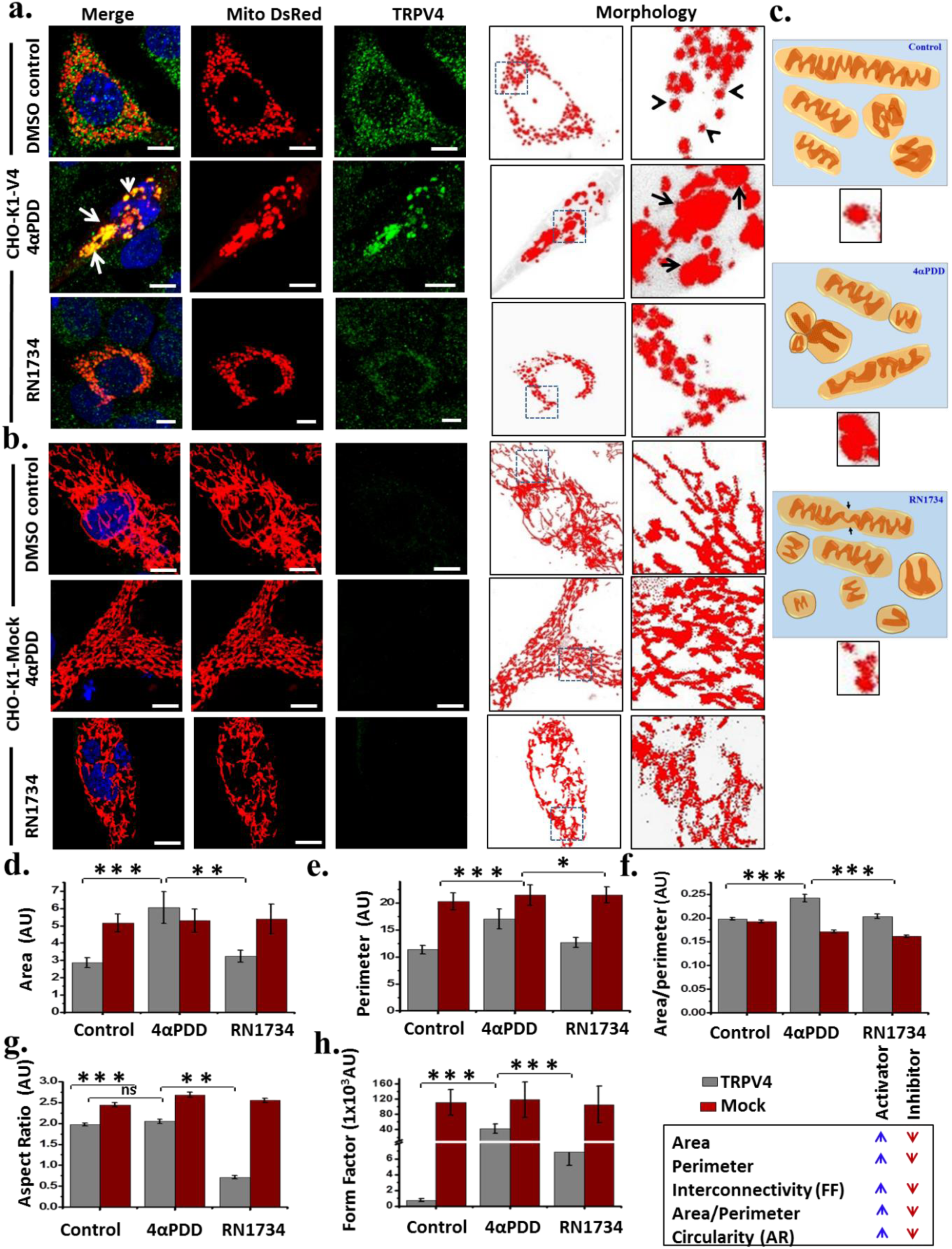
Modulation of TRPV4 alters mitochondrial morphology. a-c. CHOK1-V4 (**a**) or in CHOK1-Mock cells (**b**) expressing MitoDsRed was used and TRPV4 was activated with 4αPDD (5µM) or inhibited by RN1734 (10µM) for 8 hours. Mitochondria in CHOK1-V4 cells become spherical or round ball-like in shape after activation with 4αPDD (shown as arrow head) as compared to cylindrical or rod-like normal mitochondria observed in CHOK1-Mock cell (lower panel). TRPV4 activation leads to mitochondrial aggregation in TRPV4-positive mitochondria as compared to other conditions. The digitalized image of mitoDsRed intensity and an enlarged view filed of the same are represented on the right hand side. Arrowheads and arrows are indicating the round/spherical mitochondria and aggregated mitochondria respectively. Scale bar is 5 μm. **c.** The schematic model represents the abnormalities in the mitochondrial number and morphology. In case of TRPV4 activation, several mitochondria fuse together and form aggregated mitochondria as compared to other conditions. **d-h.** Quantitative changes in mitochondrial morphology in response to TRPV4 activation or inhibition is represented. For quantitative analysis, images of mitoDsRed expressing CHOK1-V4 or CHOK1-Mock cells were processed and all mitochondrial parameters were calculated. Represented graphs depict mitochondrial area (d), perimeter (e), area/perimeter (f), which increases significantly after TRPV4 activation in CHOK1-V4 cells. Aspect Ratio (AR) of mitochondria remain unchanged in presence of TRPV4 agonist in CHOK1-V4 cells (g). However, in CHOK1-Mock cells mitochondrial AR is higher (indicating elongated mitochondria) as compared to TRPV4-positive cell. The Form Factor (h) of TRPV4-postive mitochondria is 1000 fold higher in presence of 4αPDD as compared to control indicating that in 4αPDD treated condition mitochondria aggregates and gets interconnected to each other.

To confirm such changes in a more quantitative manner, several parameters were considered and more than 500 individual mitochondria were quantified for each parameter (Fig 3d-f). Results indicate that the mitochondrial area is higher in the presence of TRPV4 activator as compared to inhibitor in CHOK1-V4 cells. It was observed that Aspect Ratio (AR; Major axis/Minor axis) of CHOK1-V4 mitochondria is not significantly different in presence of its activator but it decreased in presence of inhibitors. However, the AR change was significant with respect to mitochondria from CHOK1-Mock cell even in the control conditions suggesting that presence of TRPV4 is sufficient to bring certain changes in the mitochondrial morphology (Fig 3g). “Aspect Ratio” is a reliable indicator of mitochondrial length; and therefore the results indicate that in CHOK1-V4 cell, the mitochondria is spherical in nature as compared to cylindrical or tubular mitochondria present in CHOK1-Mock cell. Similarly, mitochondrial “Form Factor” (FF, Pm2/4πAm) represents the branching or interconnectivity of mitochondria relevant for mitochondrial mass exchange. This FF is significantly higher (1000 fold) in TRPV4 activator treated mitochondria as compared to control and RN1734 treated mitochondria (Fig 3h). These results indicate that in the presence of 4αPDD, multiple spherical-shaped mitochondria fuse with each other to form big aggregated mitochondria (Represented as model images in Fig 3c). Taken together, these results indicate that TRPV4 modulates mitochondrial morphology, at least in stable cell lines expressing TRPV4.

The next aim was to explore if endogenous TRPV4 regulates mitochondrial morphology in a similar manner. For that purpose, we used HUVEC primary cells; as TRPV4 is reported to be expressed endogenously by these cells (Ma et al., 2010). Immunoblot and immunofluorescence analyses confirmed the endogenous TRPV4 expression (Supplementary Fig 4a). HUVEC cells were cultured and treated with TRPV4 activator or inhibitor. To explore the changes in mitochondrial morphology, treated cells were immunostained for Hsp60. In presence of TRPV4 activator 4αPPD (5 µM) mitochondria aggregate in the perinuclear area (indicated by arrows in Supplementary Fig 4b). This effect was not observed in cells that were treated with RN1734 (20 µM). In this condition mitochondrial morphology was tubular and branched similar to DMSO control. These results indicate that activation of endogenous TRPV4 is also able to induce changes in mitochondrial structure and morphology. This suggests that the regulation of mitochondria by TRPV4 may be a common phenomenon relevant in different types of cells and tissues.

### The C-terminus of TRPV4 is sufficient for mitochondrial localization

Next attempt was taken to decipher which segment of TRPV4 is really imported inside mitochondria. To understand this, TRPV4 deletion constructs [N-terminus (1-465aa), C-terminus (718-871aa) and full TM region (466-711aa)] were cloned into RFP and/or in GFP vectors (Fig 4a). Subsequently transient transfection was performed in HaCaT cells. Co-localization experiments indicate that both N-terminal fragment of TRPV4 (TRPV4-Nt-RFP) and the TM fragment of TRPV4 (TRPV4-TM-GFP) get distributed diffusely throughout the cytoplasm. Merged images of TRPV4-TM-GFP and TRPV4-Nt-RFP do not show any co-localization with immunostained Hsp60 (Fig 4b) or Mito-GFP. Cells displaying higher expression of TRPV4-Ct-RFP appear as larger aggregates and eventually die within 12 hours suggesting that over expression of TRPV4-Ct-RFP is deleterious in nature. Nevertheless, TRPV4-Ct-RFP perfectly co-localizes with Mito-GFP in cells suggesting that such larger aggregates can be actually aggregated mitochondria (data not shown).

**Figure 4.**
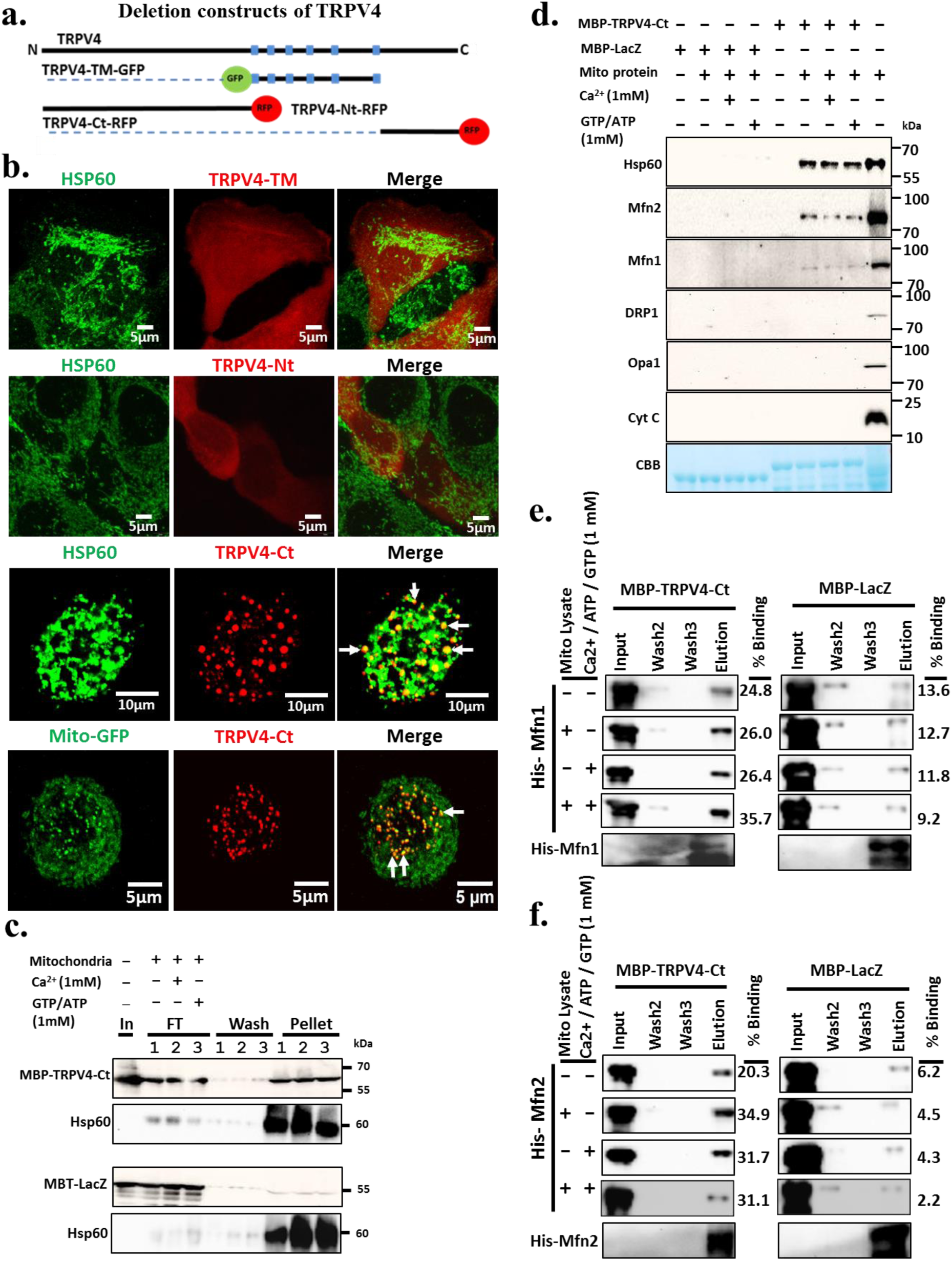
The C-terminus of TRPV4 interacts with different mitochondrial proteins. **a.** Schematic diagram representing the deletion constructs of TRPV4 used for the co-localization study. **b.** Confocal images representing the localization of TRPV4-Nt-RFP, TRPV4-TM-GFP and TRPV4-Ct-RFP with respect to mitochondrial markers. **c.** Equal amounts of purified MBP-TRPV4-Ct and MBP-LacZ was used for binding assay with intact mitochondria in presence or absence of Ca^2+^ and GTP/ATP. Eluted protein samples were subsequently probed with anti-MBP and Hsp60 antibodies. **d.** Purified MBP-TRPV4-Ct or MBP-LacZ was used for pull down experiments with mitochondrial lysate in presence of Ca^2+^ and GTP/ATP independently. Western blot analysis indicates that MBP-TRPV4-Ct, but not MBP-LacZ interacts with Hsp60, Mfn2, Mfn1 independently of the presence of Ca^2+^ and GTP/ATP. Other mitochondrial proteins such as Opa1, DRP1 and Cyt C are not detectable among the eluted proteins. Presence of MBP-TRPV4-Ct or MBP-LacZ at equal amounts in the final eluted fractions is demonstrated by Coomassie staining. **e-f.** His-Mfn1 and His-Mfn2 interact directly with MBP-TRPV4-Ct but not with MBP-lacZ. Purified His-Mfn1 or His-Mfn2 immobilized on Ni-NTA beads and purified MBP-TRPV4-Ct or MBP-LacZ were incubated under various conditions as indicated and analysed by immunoblot analyses. The values mentioned in the right side represents the percentage [with respect to the input] of protein eluted. (In: Input amount of MBP-TRPV4-Ct/MBP-LacZ, FT: Supernatant collected after incubation, W2: 2nd wash fraction, Mito: Mitochondrial pellet fraction).

To confirm that TRPV4-Ct-RFP is indeed localizing to mitochondria, we explored the co-localization between TRPV4-Ct-RFP and Mito-GFP in HaCaT cells, which express both but at very low levels and at early time points (3-6 hours after transfection). In such conditions, we observed that the C-terminus of TRPV4 appear as big dots, which are spherical in shape. These dots are distributed throughout the cytoplasm and such dots perfectly co-localize with Mito-GFP (Fig 4b). This suggests that these TRPV4-Ct-RFP positive structures can be spherical-shaped mitochondria. It also indicates that localization of TRPV4-Ct inside mitochondria significantly alters mitochondrial morphology. To confirm that such localization in mitochondria is not due to over-expression, HaCaT cells expressing low level of TRPV4-Ct-RFP were immunostained with Hsp60. Significant co-localization is observed between TRPV4 and Hsp60 in cells expressing low levels of TRPV4-Ct-RFP (Fig 4b).

### The C-terminus of TRPV4 binds with intact mitochondria and interacts with Hsp60, Mfn1 and Mfn2

As TRPV4-Ct-RFP is sufficient to localize within mitochondria, further attempt was taken to characterize if TRPV4-Ct associates with intact mitochondria in *in vitro* conditions. To this end, binding experiments were performed using purified MBP-TRPV4-Ct or MBP-LacZ (as a negative control) with intact mitochondria freshly isolated from Goat brain. MBP-TRPV4-Ct or MBP-LacZ was incubated in mitochondrial isolating buffer (in order to maintain osmolarity of mitochondria during incubation) followed by washing of unbound protein by centrifugation. Results suggest that MBP-TRPV4-Ct but not MBP-LacZ associates with intact mitochondria (Fig 4c). Next we attempted to identify mitochondrial proteins, which can potentially interact with TRPV4-Ct. Pull down experiments were performed with MBP-TRPV4-Ct or with MBP-LacZ with goat brain mitochondrial lysate alone or supplemented with Ca^2+^, GTP and ATP. Immunoblot results indicate that Hsp60, Mfn1 and Mfn2 interact with MBP-TRPV4-Ct. These interactions remain unaltered in the presence or absence of Ca^2+^ and ATP/GTP (1 mM each). However, immunoblot with anti-Opa1, anti-Cyt C and anti-DRP1 antibodies, failed to detect any of these proteins in the pull down samples (Fig 4d). These results suggest that the MBP-TRPV4-Ct indeed interact with Hsp60, Mfn1 and Mfn2 present in mitochondrial lysates.

### The C-terminus of TRPV4 interacts directly with mitochondrial dynamics regulatory proteins Mfn1 and Mfn2 independent of Ca^2^+ and GTP

Next we sought to confirm if the interaction of MBP-TRPV4-Ct with mitochondrial fusion regulatory proteins Mfn1 and Mfn2 is direct and independent of any other proteins/factors present in the mitochondrial lysate. For that purpose, His-Mfn1 and His-Mfn2 were expressed and purified. Interaction with these purified proteins (Mfn1 and Mfn2) was analysed in the presence of very less amount of mitochondrial lysate and/or combination of Ca^2+^/ATP/GTP. After interaction and substantial washing with 20 mM imidazole; all interacting proteins were eluted in 100 mM imidazole and subjected for SDS-PAGE. To visualize proteins interacting with MBP-TRPV4-Ct, western blot was performed with anti-MBP antibody. These results indicate that both His-Mfn1 and His-Mfn2 interacts directly with MBP-TRPV4-Ct but not with MBP-LacZ. This interaction is independent of the presence of mitochondrial lysate, Ca^2+^, ATP and GTP (Fig 4e-f).

### TRPV4 regulates mitochondrial membrane potential

Under normal conditions mitochondrial inner membrane is impermeable to any ions and influx of Ca^2+^ into mitochondria is controlled by various Ca^2+^-uniporters or exchangers. It has been reported that excess Ca^2+^-influx into mitochondria results in collapse of mitochondrial membrane potential (Talbot et al. 2007; Chalmers et al. 2008). In this context, it is important to explore if TRPV4 activation can cause changes in the mitochondrial membrane potential. For this purpose CHOK1-V4 and CHOK1-Mock cells were treated with TRPV4 activator or inhibitor and subsequently stained with ratiometric dye JC-1. In the presence of TRPV4 activator 4αPDD (5μM), mitochondrial potential was significantly decreased in CHOK1-V4 cells as compared to DMSO control. TRPV4 inhibitor, RN1734 (10 μM) did not alter the mitochondrial potential in CHOK1-V4 and retains its membrane potential similar to the control condition. No effect of TRPV4 activator or inhibitor was observed in CHOK1-Mock cell (Fig 5a). Subsequently image intensity (Red and Green) was quantified by Image J. The Red intensity (Ex535/Em590) images representing higher membrane potential is shown at the right side (Fig 5a). CCCP (5 μM), an uncoupler commonly used to reduce mitochondrial oxidative potential, was used as a positive control in this experiment. Quantitative results indicate that mitochondrial membrane potential in control set of CHOK1-V4 cell is significantly lower as compared to the CHOK1-Mock cells (Fig 5b). Interestingly, effects of TRPV4 activator (4αPDD) on mitochondrial membrane potential in CHOK1-V4 cells are comparable to the CCCP treated cells. Taken together these results suggest that TRPV4 may act as a mitochondrial uncoupler.

**Figure 5.**
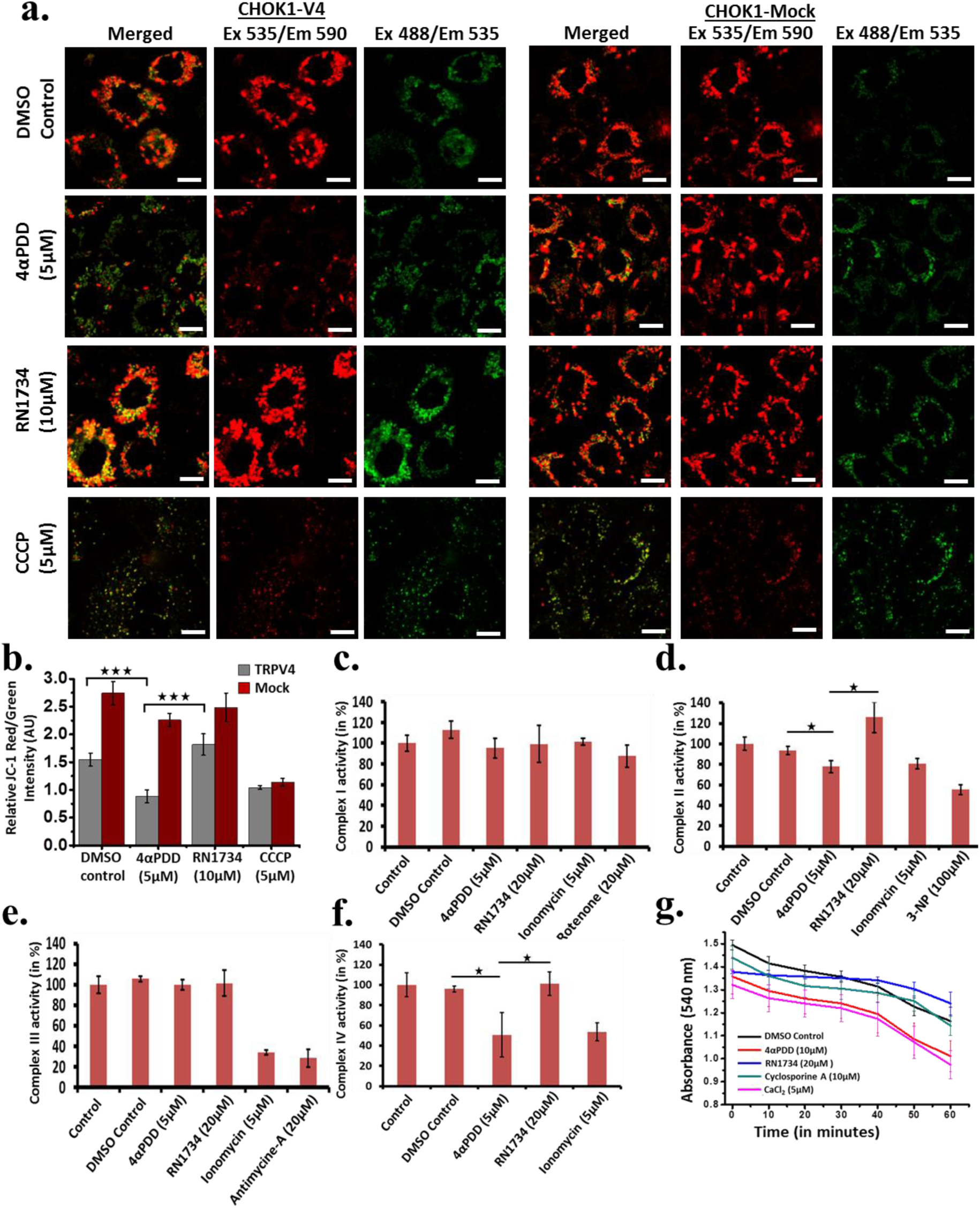
TRPV4 regulates mitochondrial membrane potential and Electron Transport Chain functions. **a.** CHOK1-V4 and CHOK1-Mock cells were treated with TRPV4 activator 4αPDD (5 μM) and inhibitor RN1734 (10 μM) for 8 hours. Subsequently JC-1 (5 μM) was added and confocal images were acquired by dual excitation wavelength 488nm (Green, for mitochondria with formation of JC-1 monomers at low mitochondrial potential) and 535nm (Red, for mitochondria with formation of J-aggregates at high membrane potentials). Scale bar is 10 μm. **b.** Red/Green intensity (in arbitrary unit) of more than 15 view fields for each condition was calculated by ImageJ. TRPV4 activation significantly decreases the mitochondrial membrane potential as compared to the control. Basal level mitochondrial potential is higher in CHOK1-Mock cell as compared to CHOK1-V4 cells. Statistical paired two test was performed and P-values are significant (n = 3). Bar graph representing the ±SEM. **c-f.** TRPV4 activation alters mitochondrial (ETC). Enzymatic activity of mitochondrial Electron Transport Chain complex I, II, III and IV was determined in isolated mitochondria purified from brain. Mitochondria were pre incubated with TRPV4 activators or inhibitors and the samples were analysed for enzymatic activity. Results indicate that enzymatic activities of Complex I and III are not altered significantly in presence of TRPV4 activator or inhibitor (c & e). However enzymatic activities of complex II and IV is significantly altered due to TRPV4 activation or inhibition (d & f). In each case, Ionomycin (Ca^2+^ ionophore) and complex chain inhibitor shows significant decrease in complex activity. **g.** TRPV4 regulates Membrane Permeability Transition (MPT) in isolated mitochondria. TRPV4 activation by 4αPDD induces more MPT as compared to control. Similarly, TRPV4 inhibition by RN1734 reduces the formation of MPT inside the mitochondria. CaCl2 is used as a positive control for MPT.

### TRPV4 alters mitochondrial Electron Transport Chain (ETC) activity and forms Mitochondrial Transition Pore (MTP)

To explore the Role of TRPV4 activation or inhibition on the activities of enzymes involved in mitochondrial electron transport chain, mitochondria were freshly isolated from goat brain and mitochondrial complex activity assay were performed (Fig 5c-f). TRPV4 activator or inhibitor was added to isolated mitochondria and enzymatic activities were assayed against different mitochondrial Complexes (I-IV). This analysis indicates that neither activation nor inhibition altered the Complex I and Complex III activity (NADH: Ubiquinone oxidoreductase and CoQ Cyt C oxidoreductase) significantly. Addition of Complex I-specific inhibitor (Rotenone) and Complex III-specific inhibitor (Antimycine-A) shows significant decrease in the activities of these complexes with respect to the control conditions. However, activity of Complex II decreased significantly in presence of TRPV4 activator 4αPDD (5 μM). However in presence of TRPV4 inhibitor RN1734 (20 μM), Complex II activity is significantly increased with comparison to control and 4αPDD (5 μM). Calcium ionophore, namely Ionomycin and complex II inhibitor 3-NP (100 μM) also cause decreased activity of Complex II. The Complex IV activity was also measured in similar manner. In presence TRPV4 activator, 4αPDD (5 μM), the Complex IV activity was decreased significantly. In contrast, presence of TRPV4 inhibitor RN1734 (20 μM) increases the complex IV activity significantly as compared to 4αPDD treated sample. In presence of Ionomycin, Complex IV activity is significantly lower as compared to other conditions. Mitochondrial Complex activity assays suggest that TRPV4 activator or inhibitor largely regulate the function of Complex II and IV activity as comparison to the Complex I and III.

Mitochondria forms large conducting pores in the inner mitochondrial membrane in case of over saturation of Ca^2+^ ions and this is known as Membrane Permeability Transition (MPT). To know the effect of TRPV4 activator or inhibitor upon the formation of mitochondrial membrane pore (Mitochondrial swelling), we performed MPT assay with mitochondria freshly isolated from goat brain. MPT results indicate that in presence of TRPV4 activator 4αPDD (10 μM), mitochondria get swollen up and form MPT and its absorbance decreases with time (Fig 5g). However in presence of TRPV4 inhibitor RN1734 (20 μM), the mitochondria did not form MPT as compared to others. Addition of CaCl_2_ (1 mM, a known inducer of MPT) in mitochondria was taken as a positive control. The MPT graph of 4αPDD shows similar changes like CaCl_2_ in later time point which suggest that TRPV4 activator regulates MPT in isolated mitochondria.

### TRPV4 regulates mitochondrial Ca^2^+-dynamics

Ca^2+^-influx inside the mitochondria occurs through outer mitochondrial membrane followed by inner membrane. Outer mitochondrial membrane allows free transport of Ca^2+^ ions but the inner membrane is impermeable for Ca^2+^. VDAC and many other unidentified ion channels are present on the outer membrane of mitochondria which allow passing of cytoplasmic Ca^2+^ ions across the outer mitochondrial membrane. As TRPV4 is present in mitochondria, we explored if activation of TRPV4 can cause Ca^2+^-influx in mitochondria. To explore the mitochondrial Ca^2+^ dynamics in presence of TRPV4 activator or inhibitor, live cell imaging was performed in CHOK1-V4 and CHOK1-Mock cell. For this purpose, Mito-pericam (Mitochondrial Ca^2+^-sensing construct) was transiently transfected in CHOK1-V4 and CHOK1-Mock cell and live cell imaging was performed. Cells exhibiting moderate levels of expression were considered for Ca^2+^ imaging because these cells allow qualitative analysis of both increase as well as decrease in the intensities. This experiment suggests that Ca^2+^-influx inside mitochondria increases after addition of TRPV4 activator 4αPDD in CHOK1-V4 cells (Fig 6a & c). Addition of TRPV4 inhibitor RN1734 results in decrease in mitochondrial Ca^2+^ level (as indicted by fluorescence intensities of Mito-pericam) with progress in time. However, Ca^2+^-influx in CHOK1-Mock cell was not altered much in presence of TRPV4 activator or inhibitor (Fig 6b & d).

**Figure 6.**
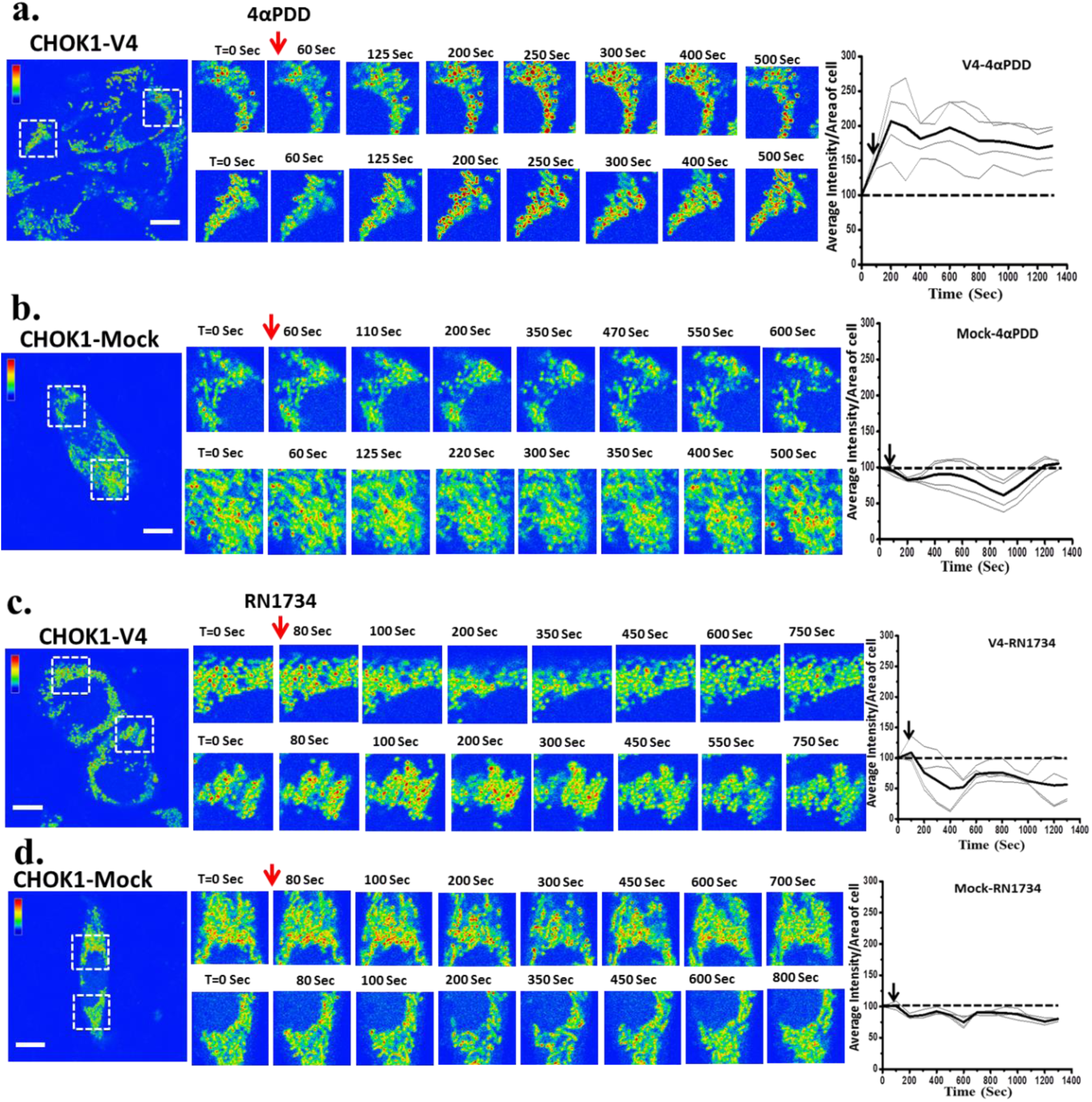
TRPV4 regulates mitochondrial Ca^2+^-influx. Mito-pericam was transiently expressed in CHOK1-V4 or CHOK1-Mock cells and live cell imaging was performed to monitor the effect of TRPV4 activation (4αPDD, 5 μM) **(a-b)** or inhibition (RN1734, 10 μM) **(c-d).** Addition of 4αPDD causes massive mitochondrial Ca^2+^-influx in CHOK1-V4 cell (a). However addition of RN1734, reduces the mitochondrial Ca^2+^-level with time **(c)**. In CHOK1-Mock cell, Ca^2+^-levels remain mostly unchanged in the presence of TRPV4 activator **(b)** or inhibitor (d). Ca^2+^ intensity graphs of multiple alive cells in each condition was quantified by ImageJ are represented on the right side). Thick dark black line represents the average value of mitochondrial Ca^2+^-level (Arrow indicates the time of addition of drug, dotted line represents initial value as 100%). Scale bar: 20 μm.

### TRPV4 activation altered mitochondrial structure and its organization in other cellular system

So far obtained data suggests that TRPV4 may be a part of mitochondria-associated ER membrane (MAM) portion which connects ER to mitochondria and therefore contribution of ER cannot be completely ruled out. In order to prove that TRPV4 indeed can regulate mitochondrial functions, we have tested the importance of TRV4 in mature mammalian spermatozoa which is an ER-free primary cellular system. Recently we have demonstrated the conserved endogenous expression of TRPV4 in all vertebrate sperm (Kumar et al. 2016). Therefore we aimed to investigate if TRPV4 colocalizes with mitochondrial markers. Our results confirm distinct colocalization of TRPV4 with mitochondrial markers. In most of the cases, TRPV4 is enriched in mid piece region and it colocalizes with specific mitochondria (but not with all) that are labelled with MitoTracker Red (indicated by arrows) (Fig 7a). Previously we have demonstrated that TRPV4 activation alters the Ca^2+^-level in the mitochondria-enriched neck regions suggesting that mitochondrial coiling at the neck regions may get affected by TRPV4 activation (Kumar et al. 2016). To explore the changes in mitochondrial morphology and its coiling in bull sperm, TRPV4 activator 4αPDD (1μM) was added in sperm and incubated for 2 hours at 37°C. Subsequently, MitoTracker Red was added to label the mitochondria. Results indicate that in presence of TRPV4 activator, in many cases, sperm mitochondria formed blebbing or split in the mitochondrial structure (Fig 7b). Such abnormalities were not found in the control cases (n = ∼500 cells). This in general suggests that TRPV4 activation induces extreme abnormalities in the mitochondrial organization. To observe minute changes at better resolution, we performed super resolution microscopy and analysed the mitochondrial coiling pattern in control and TRPV4-activated conditions. Results suggest that in control condition, sperm mitochondrial coiling was intact and form a perfect helix-like structure with regular thickness and pitch length in the midpiece region (Fig 7c). However, in presence of TRPV4 activator, mitochondrial coiling and its organization drastically changes. Helical organization of mitochondria was altered and they formed “kink” or “small cervices-like structure” in the mitochondrial ring structure (Fig 7c). Other defects such as bending patterns were also altered to some extent (data not shown).

**Figure 7.**
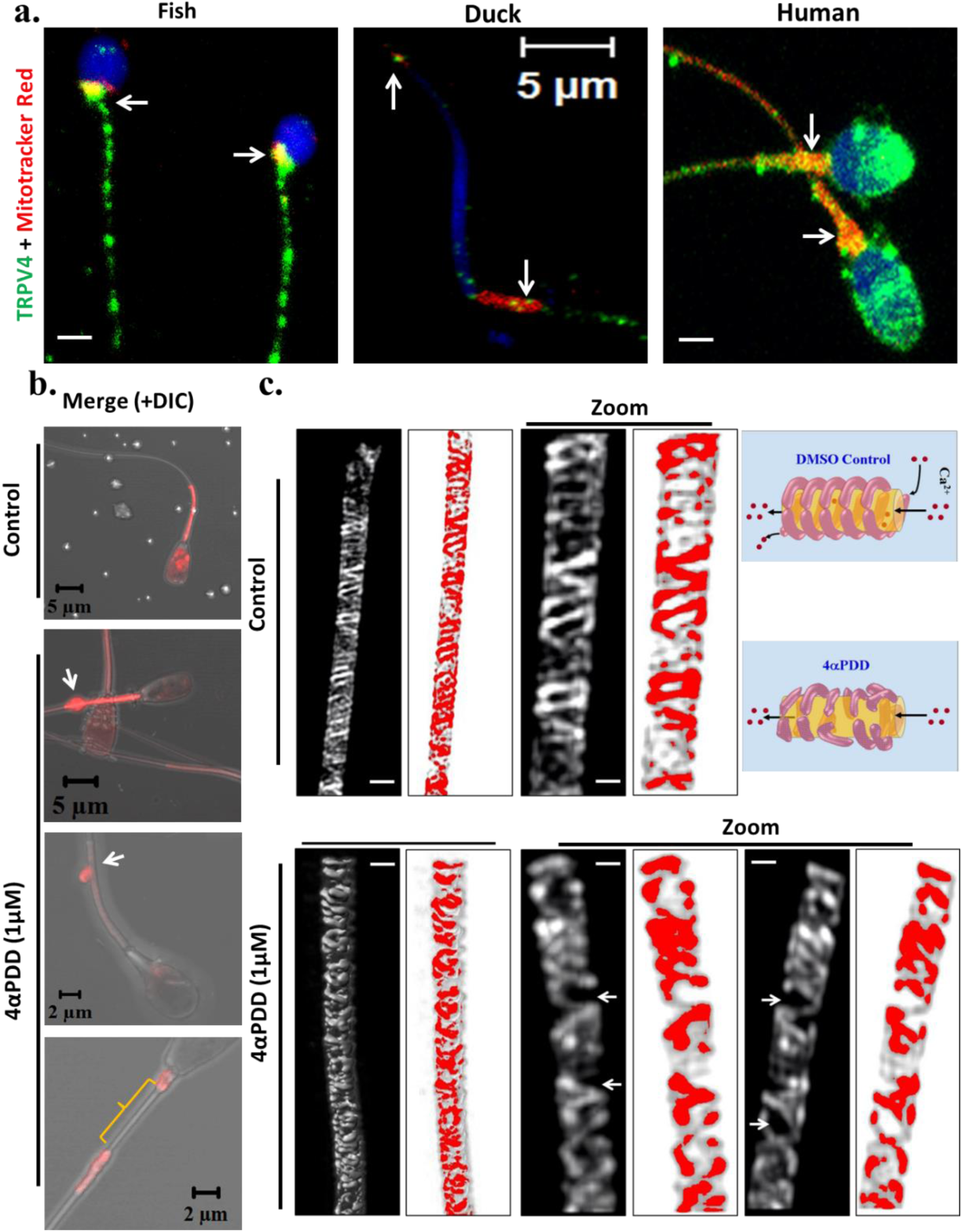
TRPV4 localizes in sperm mitochondria of different vertebrates and activation of TRPV4 disrupts mitochondrial coiling. **a.** Confocal images demonstrating the colocalization (indicated by arrows) of TRPV4 (green) with MitoTracker-Red labelled structures in fish, duck and human sperm. Some but not all sperm mitochondria revel the presence of TRPV4. In case of duck sperm, apart from mid piece region, colocalization is also observed at the tip of the head. **b.** Confocal images depicting the mitochondrial abnormality observed in bull sperm when TRPV4 is activated by 4αPDD (1 μM, 2 hours at 37°C). Activation induces blabbing or node-like structure (indicated by arrow) in the mid-piece regions. In some cases, sperm mitochondrial structure becomes fragmented into two halves. **c.** Shown are the super resolution (SIM) images demonstrating the mitochondrial coiling pattern in control and TRPV4-activated conditions. In control condition, mitochondrial coiling is intact and form regular helix-like structure. In TRPV4-activated condition, the regular helical organization is disrupted (kinks or cervices in the mitochondrial coiling region are indicated by arrows). For better visualization enlarged images are presented in each panel. Models showing the Ca^2+^-influx and abnormalities in the mitochondrial coiling in presence of TRPV4 activator. Scale bar: 5 μm (for left side panel) and 1 μm (for enlarged image).

Taken together, our results demonstrate for the first time that TRPV4 is a mitochondrial protein, interacts with intact mitochondria as well as specific mitochondrial proteins and it regulates mitochondrial structure-function, a phenomena that is mostly conserved in different cellular systems.

### Discussion

The importance of mitochondria in the intracellular Ca^2+^-buffering is well established (Budd and Nicholls. 1996; Duchen et al. 1999; Jouaville et al. 1995). In addition, for a long time it is well known that the temperature-sensing ability is a unique property of cellular mitochondria, even in isolated conditions though the molecular players involved in this feature are currently unknown (Johnston et al. 1994; Kang et al. 2008; Chrétien et al. 2018). In this work, we demonstrate that the TRPV4 channel localizes to a sub-population of mitochondria in various cell types. TRPV4-mediated mitochondrial regulation thus seems to be universal in nature and is also observed in different tissues including the mammalian brain, adipocytes and other primary cells such as HUVEC cells and in mature spermatozoa from different vertebrates. Our results shed light on important and novel aspects of TRPV4, and expand the significance of TRP channels in general.

### Presence of TRP channels in mitochondria

Though initially it was assumed that all TRP channels are primarily localized in the plasma membrane, later reports have confirmed that different TRP channels have different yet specific distributions to subcellular membrane systems, including ER, Lysosome or endosome, where they participate complex tasks such as regulation of membrane trafficking signal transduction, maintenance of ion and pH homeostasis (Dong et al. 2010; Yadav & Goswami. 2016). For example, in DRG neurons, TRPV1 is physically and functionally present in the ER membrane where it regulates the Ca^2+^-wave at distinct microdomains, what may cause Ca^2+^-overload at ER-associated mitochondria and subsequent damage of these mitochondria (Gallego-Sandín et al. 2009; Olah et al. 2001). Similarly, TRPC3 is physically and functionally present in mitochondria and regulates membrane potential as well as Ca^2+^ oscillation inside these mitochondria (Feng et al. 2013). However, the molecular identities of various Ca^2+^ channels and particularly different TRP channels present in mitochondrial membranes are still lacking. Here, we demonstrate that TRPV4, a temperature sensitive ion channel is endogenously present in mitochondria. Our work also provides molecular mechanisms that involve TRPV4 in the regulation of mitochondrial structure-function relationship. For example we show that activation of TRPV4 results in Ca^2+^-influx into mitochondria. This observation is in full agreement with our previous report demonstrating that the Ca^2+^-pattern in mitochondria-enriched neck region of mature spermatozoa is altered in case of TRPV4 activation (Kumar et al. 2016). Moreover, the presence of TRPV4 that can act as a thermo-sensitive ion channel (Guler et al., 2002), agrees well with the reports that mitochondria maintain higher temperature than the cytosol (Chrétien et al. 2018; Lane N 2018).

### TRPV4 interacts with mitochondrial proteins and is part of MAM

The observation that the C-terminus of TRPV4 interacts with key mitochondrial proteins, namely with Hsp60 and with Mitofusions gives rise to the hypothesis that TRPV4 may contribute to essential mitochondrial functions including fusion and fission events These findings are also relevant for the understanding of diseases involving mitochondrial abnormalities. To date, several pathophysiological conditions are known that are associated with mitochondrial abnormalities (Michael et al. 2006). Involvement of TRPV4 in different tissue functions is well established and mutations and/or abnormalities in TRPV4 is associated with a variety of pathophysiological conditions commonly referred to as channelopathies (Verma et al. 2010). For example, more than 20 genes including TRPV4 and Mfn2 are considered to be involved in the development of Charcot Marie Tooth 2 disease (defects in sensory/motor neurons) suggesting that TRPV4 is physiologically linked with Mfn2 (Bird TD. 1998). Indeed our experiments confirm that TRPV4 interacts with Mfn1 and Mfn2 directly and thus support the genetic evidence at the protein level.

The interaction of Mfn1 and Mfn2 with TRPV4 also suggests that TRPV4 might be present in mitochondria-associated ER membrane (MAM) fraction, the very specialized region which maintains ER-mitochondrial contact sites, i.e. the micro-domain relevant for Ca^2+^ dynamics in a range of 20-200 nm distance (Patergnani et al. 2011). This is also consistent with the observation that TRPV4-positive mitochondria are primarily localized in the perinuclear regions. MAM is closely connected to ER and thus may facilitate the direct sorting of TRPV4 from ER to mitochondria (Raturi and Simmen. 2013). Notably, Mfn2 is known to be involved in the regulation of endoplasmic reticulum-mitochondrial contact regions (Sugiura et al. 2013) and thus TRPV4-Mfn2 complex may be relevant for Ca^2+^-dependent regulation of mitochondrial functions (De Brito and Scorrano. 2008; Qi et al. 2015). In this context, it is important to mention that mitochondria have sensitivity for recognizing microdomains of high cytoplasmic Ca^2+^, which dissipates high influx and efflux of Ca^2+^ across the mitochondrial membrane. Because of this notion most of the mitochondria are in close proximity with ER (in fact connected through MAM) and mediate easy exchange of divalent cations such as Ca^2+^. Recent reports also suggest that MAM regions are relevant for several neurological disorders, as mitochondrial functions are related with critical functions, such as lipid homeostasis, cholesterol biogenesis, mitochondrial calcium signalling, mitochondrial morphology and mitochondrial dynamics (Krols et al., 2016). Our previous study demonstrated that TRPV4 has ability to interact with different sterols, sterol-precursors and their derivatives (Kumari et al. 2015). These findings support the notion that presence of TRPV4 in the MAM can be of functional importance for sterol transport between ER and mitochondria (Kumari et al. 2015). Altogether, our results suggest that TRPV4 is present in the MAM region and regulates ER-mitochondria associated functions significantly. However, the observation that TRPV4 is present in the mitochondria of mature spermatozoa (which is generally devoid of any ER membrane) of different vertebrates strongly confirms the *bona fide* presence of TRPV4 in the mitochondria along with plasma membrane (Kumar et al. 2016).

### TRPV4 regulates mitochondrial Ca^2^+-influx and mitochondrial membrane potentiality

Mitochondria are well known organelles required for buffering of intracellular Ca^2+^ via different uniporter or cation exchangers. Using a specific mitochondrial Ca^2+^-sensor, our results confirm that TRPV4 agonist and antagonists play a critical role in the regulation of mitochondrial Ca^2+^-levels. They also affect mitochondrial structure, membrane potential and its energetics. The increment in the mitochondrial Ca^2+^-levels after 4αPDD treatments may be mediated by TRPV4 being present in both mitochondria as well as in plasma membrane. However, similar effects of 4αPDD and CCCP in CHOK1-V4 cell lines strongly argue for a functional contribution of TRPV4 in mitochondrial membranes causing increased Ca^2+^-influx into mitochondria. Excess of Ca^2+^ in mitochondria may regulate the energetics in several manners: first, initiation of dissipation of mitochondrial membrane potential; second, induction of changes in mitochondrial morphology; third; alteration of mitochondrial metabolite exchange, ATP production and/or ETCs; and fourth, MPT is induced through which excess Ca^2+^ can be removed.

The effect of TRPV4 activator or inhibitor on mitochondrial structure and Ca^2+^-influx seems to be ubiquitous in nature. Recently we have shown that TRPV4 is present in human spermatozoa and regulates the mitochondrial calcium wave in the presence of TRPV4 activator or inhibitor. TRPV4 inhibitor restricts the calcium wave propagation through coiled mitochondria (Kumar et al. 2016). Using spermatozoa as a model system (Fish, duck and human) localization studies indicate that TRPV4 is not only present in sperm membrane but also in coiled mitochondria where it may regulate sperm Ca^2+^-wave (Kumar et al. 2016). Our present data suggest that TRPV4 activator significantly alter the mitochondrial coiling pattern in bull sperm (Fig 7c), where TRPV4 is also expressed thought at somewhat lower level. It seems that TRPV4 activator increases the fusion or interconnectivity of coiled mitochondria in sperm cells. These results fit well with the results obtained from CHOK1-V4 stable cells or in HUVEC cells (Fig 3, Supplementary fig 4).

The effect of TRPV4 activation or inhibition on mitochondrial structure and morphology has been reported previously. It is well known that mitochondrial dysfunction in neuronal cells (or other cells) may cause the production of Reactive Oxygen Species (ROS), Nitric Oxide (NO) and excesses of Ca^2+^-influx, what in turn leads to the generation of neuropathic and chronic pain (Kim et al. 2009). It seems plausible that due to TRPV4 activation, the Ca^2+^ levels increase in TRPV4 containing mitochondria, what in turn may increase the production of ROS and NO and ultimately results in neuropathic or chronic pain. An *in vivo* study also revealed that in case of neuropathic or inflammatory pain conditions, several parameters such as mitochondrial position, number and its morphology are altered (Guo et al. 2013). It was observed in the same study that mitochondrial aggregation or clustering in the perinuclear region is increased, especially in the presence of inflammatory molecules inducing neuropathic pain. Our studies strongly suggest that TRPV4 is physically present in a subset of mitochondria located in the perinuclear region and contributes significantly to the regulation of mitochondrial structure-function relationship. Thus, based on our data, a role of TRPV4 in mitochondrial abnormality, relevant in neuropathic pain and several other pathophysiological conditions can be hypothesized.

### Molecular size of TRPV4 within mitochondria

Using highly specific anti-TRPV4 antibodies, our immunoblot and immunofluorescence analysis confirms the presence of TRPV4 in the mitochondrial fraction isolated from rat brain as well as from CHOK1-V4 cells that express TRPV4 at low levels. Immunoblot analysis also confirms the endogenous presence of TRPV4 in isolated mitochondria obtained from brain tissue of rat, mice and also goat. Moreover, TRPV4 is present in the mitochondria isolated from goat adipose tissue and the synaptosomal protein fraction isolated from rat brain. In most cases, TRPV4-specific band was observed in mitochondrial fraction and the molecular size matches well with the expected molecular weight (∼98 kDa). In some cases, using an antibody specific for the C-terminal cytoplasmic domain of TRPV4, we detected specific immunoreactivity at ∼72 kDa in addition to the 98 kDa band in mitochondrial fractions isolated from several biological sources (such as in synaptosomal fraction and mitochondria isolated from goat brain). This matches well with a recent study where a TRPV4-specific band of ∼70 kDa size has been detected by Western blot analysis (Zhao et al. 2012). In the present study, we also detected TRPV4-specific band at ∼98kDa as well as another strong band at ∼72 kDa in the light membrane and in synaptosomal protein fractions. The 72 kDa band was not visible in any other fractions (e.g. in post synaptic density). In all cases, the recognition of the 72 kDa polypeptide is completely suppressed in the presence of a specific blocking peptide, suggesting that this band indeed represents TRPV4.

This smaller band of TRPV4 (∼70 kDa) present in mitochondria needs further investigation. It may represent a splice variant or a very specific truncated product formed due to specific proteolytic action within mitochondria. Further detailed studies are needed to understand the mitochondrial localization of TRPV4 and importance of different TRPV4 mutants in the context of mitochondrial functions.

## Acknowledgements

Funding from NISER and DBT (Govt. India, grant number BT-BRB-TF-2-2011 & BT/PR8004/MED/30/988/2013) are acknowledged. The funders had no role in study design, data collection and analysis, decision to publish, or preparation of the manuscript. We thank the FSB Cuttack for their support with the bull sperm samples. Help from all the lab members is appreciated. CG acknowledge the previous supports from MPI, Berlin. Financial support from NISER is appreciated. Imaging facility from NISER is appreciated. 3DSIM imaging was performed at the Dundee Imaging Facility, University of Dundee and supported by an MRC Next Generation Optical Microscopy Award (Ref: MR/K015869/1). We thank Markus Posch and Jason Swedlow access to the Dundee OMX and for recording images.

## Supplementary figure

**Supplementary Fig 1:**
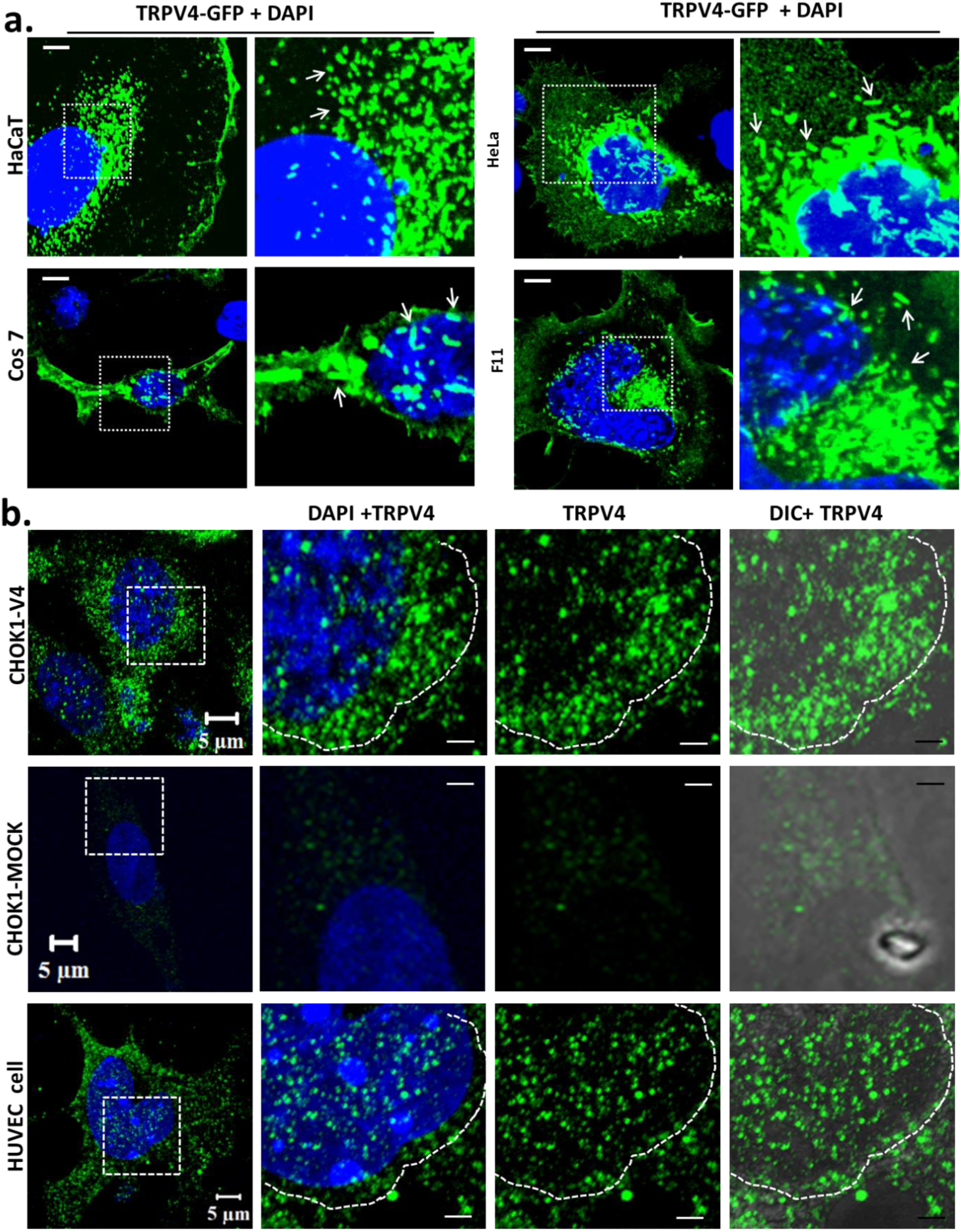
Perinuclear localization of TRPV4 in different cell lines. **(a)** TRPV4-GFP was transiently expressed in neuronal (F11) and non-neuronal (HaCaT, Cos7, HeLa) cell lines where TRPV4-specific clusters are present in the perinuclear regions. Enlarged confocal images of each are shown in the right side. **(b)** Confocal images showing that localization of TRPV4 in cells either stably expressing TRPV4 (named CHOK1-V4) or an empty plasmid (CHOK1-Mock). Endogenous expression of TRPV4 is shown in HUVEC cells (lowermost panel). TRPV4 is detected by immunostaining with TRPV4-specific antibody (green) and the nucleus is stained by DAPI (blue). Scale bar: 10 µm (a), 5 µm (b).

**Supplementary Fig 2.**
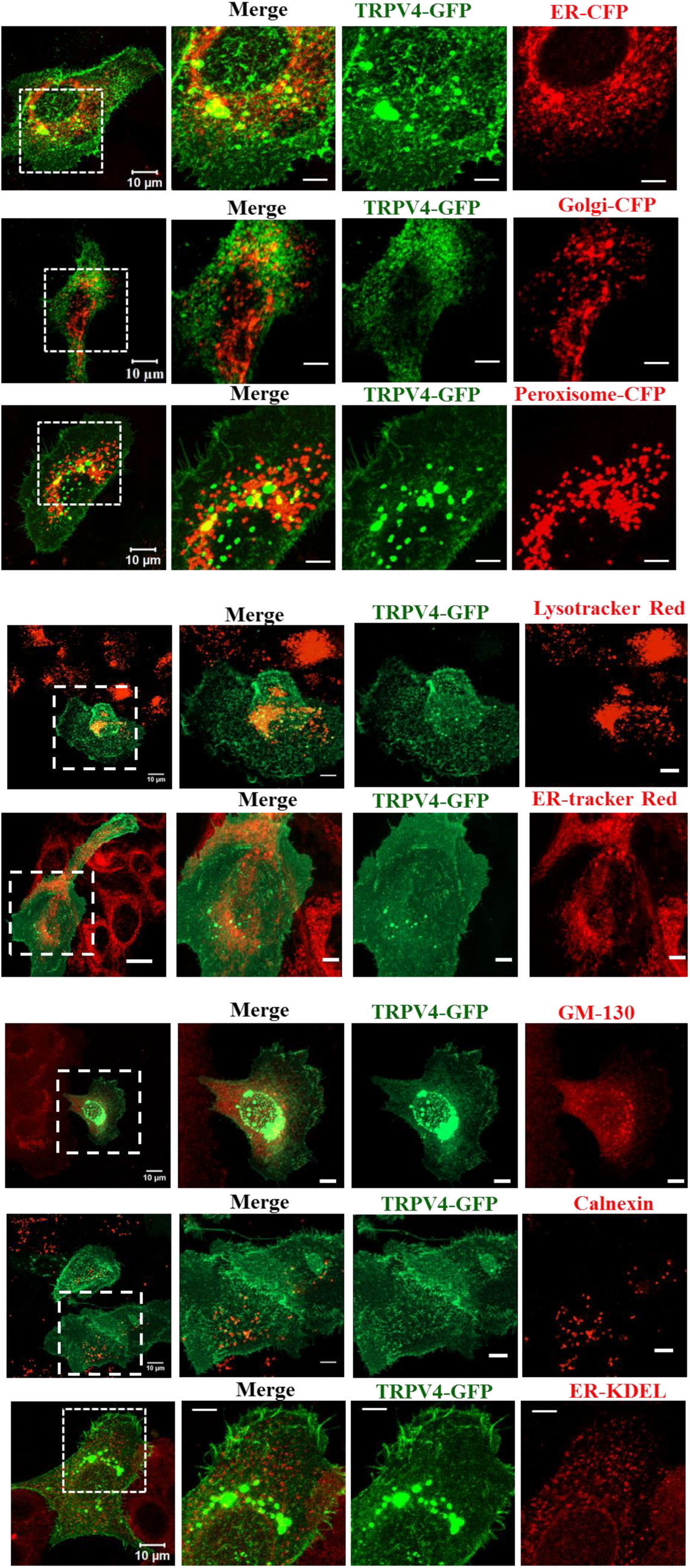
TRPV4 does not colocalizes with other subcellular organelles. Confocal images of HaCaT cells transiently expressing TRPV4-GFP and sub-cellular marker proteins such as ER-CFP, Golgi-CFP and Peroxisome-CFP are shown. Cells were also labelled with Lysotracker Red and ER-tracker are shown. Similarly, HaCaT cells expressing TRPV4-GFP are immuno-labelled for sub-cellular organelles specific markers, such as with anti GM130 ab, anti-Calnexin ab and anti-KDEL ab, Enlarged images (indicated by white dotted line) represent the perinuclear region in details. Scale bar: 5 µm (for enlarged image).

**Supplementary Fig 3.**
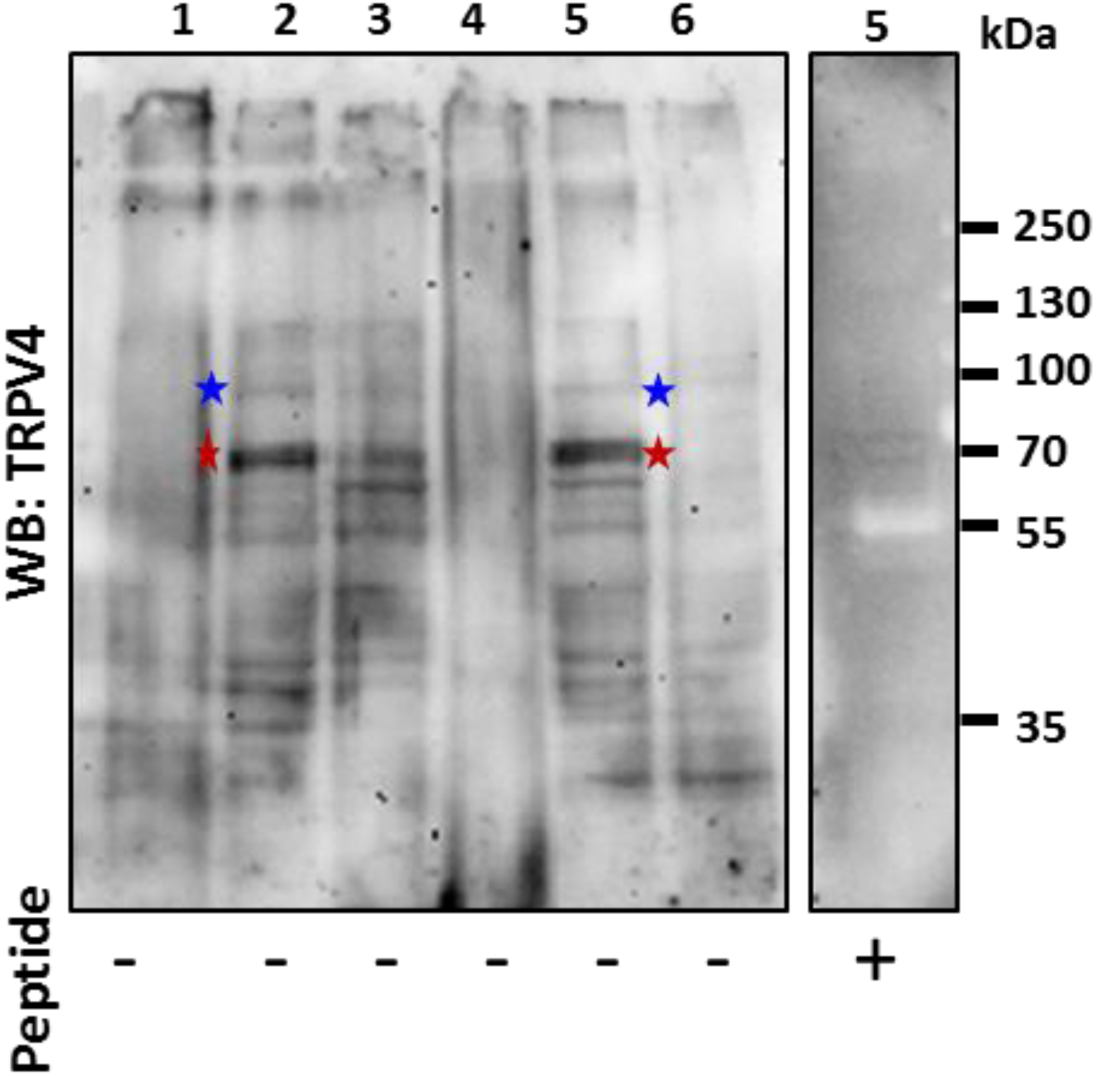
TRPV4 is present in the synaptosomal fraction of brain tissue. Tissue lysate was prepared from Rat brain and density gradient separation was done for different fractions and subsequently probed for the endogenous TRPV4. Lane 1. Microsome fraction, Lane 2: Light membrane fraction, Lane 3: Crude membrane fraction, Lane 4: Myelin fraction, Lane 5: Synaptosomal fraction, Lane 6: Synaptic junction fraction. TRPV4-specific faint band (∼ 98kDa, indicated by blue star) is present in light membrane fraction (lane 2) and in synaptosomal fraction (lane 5). In addition, another strong band for TRPV4 is also observed in these same fractions (indicated by red star). The presence of a specific peptide (corresponding to the C-terminus of TRPV4), the TRPV4-specifc band is abolished in the synaptosomal fraction (right side).

**Supplementary Fig 4.**
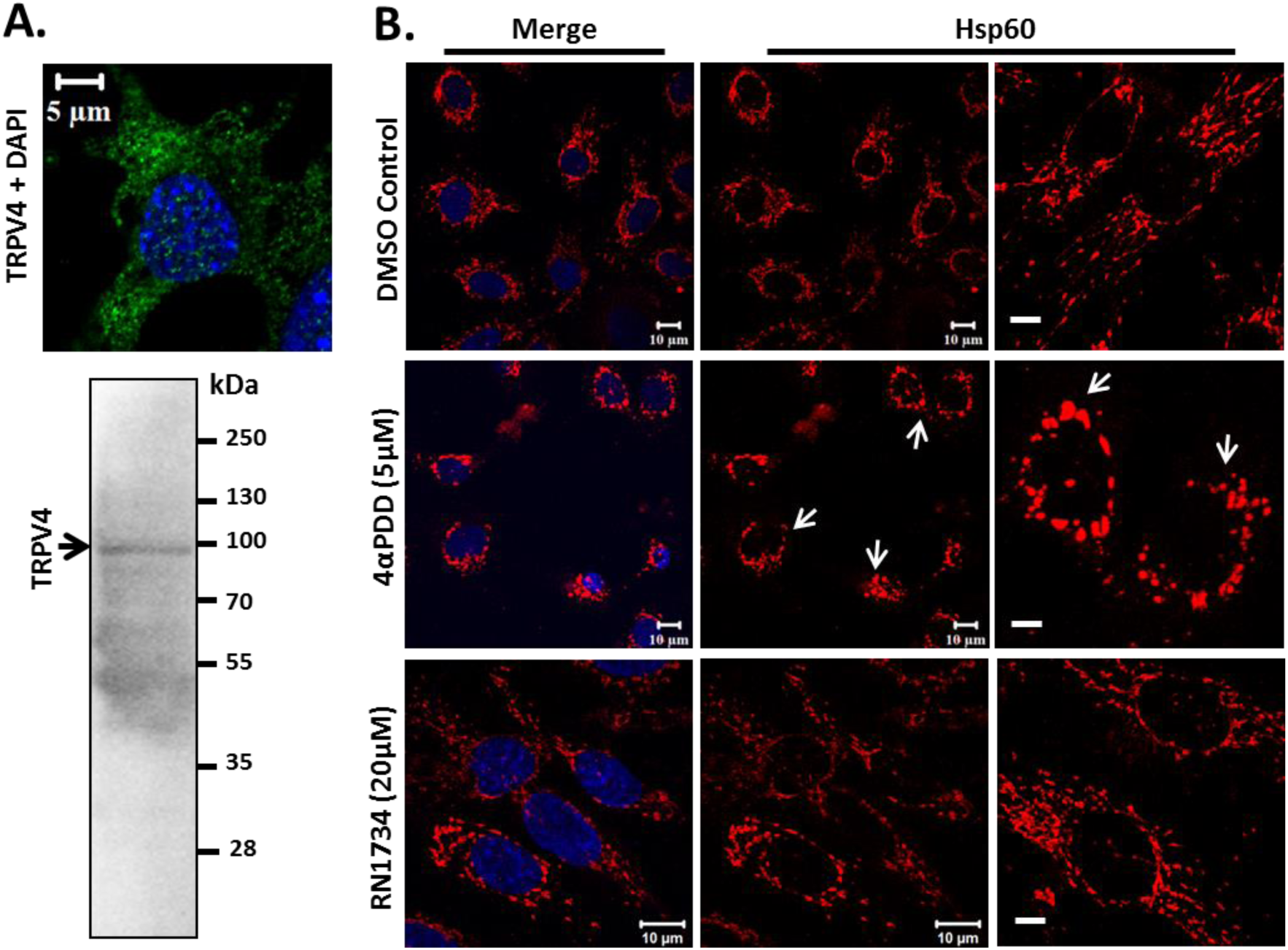
Endogenous TRPV4 regulates mitochondrial morphology in primary cell. **a.** Representative image showing the presence of endogenous TRPV4 in HUVEC cell. Western blot analysis of HUVEC cells for TRPV4 is shown below. **b.** HUVEC cells were treated with TRPV4 agonist (4αPDD) and antagonist (RN1734) and subsequently immunostained for Hsp60. Representative images depict that in presence of 4αPDD (5 µM) mitochondria become aggregated in the perinuclear region (arrows). Mitochondrial morphology was virtually normal as compared to DMSO control in the presence of inhibitor RN1734 and control. Scale bar: 5 μm.

